# Engineered OAA lectins as selective and sensitive high mannose glycan targeting tools

**DOI:** 10.64898/2026.03.04.709641

**Authors:** Bryce E Ackermann, Emerson Hall, Vanessa T Mariscal, Alex E Clark, Kevin D Corbett, Aaron Carlin, Alex J Guseman

**Affiliations:** Department of Biochemistry and Molecular Biophysics, University of California San Diego, La Jolla CA, 92039, USA; Department of Pathology, University of California San Diego, La Jolla, CA 92093, USA; Department of Molecular Biology, University of California San Diego, La Jolla, CA, 92093, USA; Department of Cellular and Molecular Medicine, University of California San Diego, La Jolla, CA, 92093, USA; Department of Medicine, University of California San Diego, La Jolla, CA 92093, USA

## Abstract

The *Oscillatoria agardhii agglutinin* (OAA) lectin interacts with N-glycans through a pentamannose core shared among all high mannose N-glycans (HMGs). Because HMGs only differ by number of mannose sugars, there is a scarcity of tools sensitive enough to resolve each specific HMG structure in their biological context. Here, we investigate the sequence space of OAA to tune the binding properties towards selectivity of Man_5_GlcNAc_2_, thus generating a structure-specific detection tool. Using phage display to screen a diverse library of OAA variants, we identify a variant with high selectivity for Man_5_GlcNAc_2_ that we further dissect to reveal four mutations necessary for selectivity and two mutations responsible for enhanced affinity for all HMGs. Coupling a crystal structure of the selective variant with binding analysis of specific point mutations, we reveal how co-dependent mutations achieve selectivity. We then demonstrate how variants can be valency-modulated on a single beta-barrel scaffold to improve their binding properties by orders of magnitude. Finally, we showcase the applicability of engineered OAA variants as improved HMG profiling tools and tunable antiviral agents.

## Introduction

Glycans are complex carbohydrates utilized by organisms to modify biomacromolecules that play key roles in protein folding, macromolecular recognition, and cell-cell signaling.^1,2^ The glycans that modify proteins are categorized as N-glycans or O-glycans, depending on whether they are attached through asparagine side chains (N-linked) or serine/threonine side chains (O-linked).^3^ All N-glycans are linked to asparagine by two N-acetylglucosamine (GlcNAc) sugars, and can be characterized into three main classes: high mannose, hybrid, and complex glycans. High mannose glycans (HMGs) are made up of only mannose sugars, other than the GlcNAc linkage (Fig. 1A). Hybrid glycans share the same pentamannose core as HMGs but have non-mannose sugars such as sialic acid and galactose extending the D1 arm. Complex glycans share only the three central mannose sugars of HMGs, and can adopt many diverse branches, lengths, and sugar varieties. N-glycans are added co-translationally as HMGs and further processed to hybrid or complex glycans in a non-templated fashion as the protein matures through the endoplasmic reticulum (ER) and Golgi. The processing of the glycans is determined by the spatial/temporal accessibility of the glycan to tailoring enzymes and is heavily influenced by cell metabolism and homeostasis. As a result, a given glycoprotein population is a heterogenous mixture of proteins with the same amino acid sequence but different glycan modifications.

**Figure 1.**
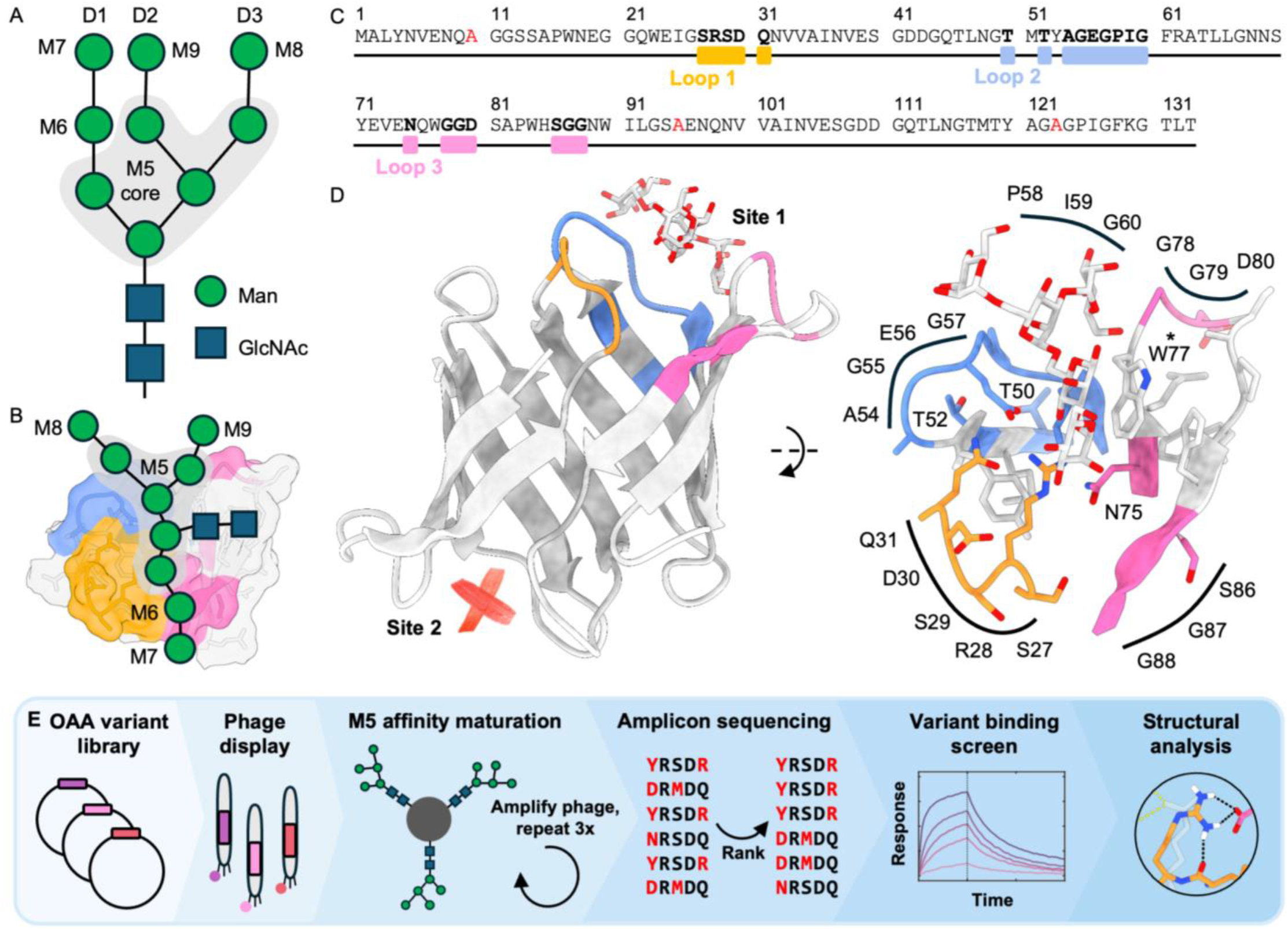
Phage display workflow and design. (A) A cartoon representation of the Man_9_GlcNAc_2_ (M9) high mannose glycan, where mannose is shown as a green circle, and GlcNAc by a blue square. The M5-core is surrounded by gray with each following mannose named as the representative glycan used in this study. Each branch of the glycan is differentiated by the nomenclature D1, D2, and D3. (B) A cartoon of the M9 glycan in the hypothetical binding pose relative to OAA. OAA is shown in surface view with loops colored according to their identity. (C) The amino acid sequence of OAA. W10, R95, and E123 are mutated to alanine (red). Loops 1, 2, and 3 are highlighted by orange, blue, and pink respectively. Residues mutated in the combinatorial library are in bold. (D) (*left)* Beta-barrel structure of OAA (PDB:3S5X)^36^ with 3α,6α-mannopentaose shown bound to site 1 with loops colored by identity. Site 2 is without ligand to represent the effect of the alanine mutations. (*right)* Close-up of site 1 with all sidechains shown. Labeled residues are colored by loop and heteroatom identity. All residues denoted here are included in the combinatorial library except for W77 marked by an asterisk. (E) A combinatorial variant library of OAA was generated semi-synthetically and cloned into pComb3xss for phage display. Phages were screened over M5-bound beads, eluted and reamplified three times. Each round of screening was assessed by amplicon sequencing and highly enriched variants were selected for binding analysis by BLI. The top hit was assessed by x-ray crystallography.

Improved understanding about the role of glycans in both cell homeostasis and disease necessitates the development of antibody-like tools to detect glycosylation. Lectins are a class of carbohydrate-binding proteins (CBPs) found in every organism in the tree of life to read out diverse glycan structures. Lectins in mammals play key roles in protein folding, signaling,^4^ and tissue self-recognition.^5^ Outside of mammalian biology, lectins are abundantly employed by plants, bacteria, and fungi as defensive detectors of non-host glycoproteins.^6,7^ Lectins themselves are similarly diverse, adopting binding pockets sensitive to minor differences between glycan sugars, linkages, and conformations.^8^ Therefore, lectins are becoming promising biotechnology candidates for anticancer drugs,^9,10^ broad-spectrum antivirals,^11,12^ immune system modulators,^13–15^ disease sensors,^16,17^ and molecular probes for glycans.^18–20^

The engineering of novel lectins and other CBPs has had relatively limited success converting into usable therapeutics or highly specific biosensors.^21–28^ Many lectins have high specificity for small oligosaccharide motifs that exist as parts of larger, more extended glycans.^29^ As a result, they may bind a high number of similar glycan structures. Additionally, most lectins are multivalent, complicating the independent evolution of binding sites. While these features may enhance lectin function in their native context, such promiscuity can lead to off-target effects on normal cells^13,30^ and non-specific analytics,^29^ limiting the scope of lectin applications. Thus, to evolve lectins to have desired properties we must choose a scaffold that i.) interacts with key regions of the glycan; ii.) is stable upon mutation; and iii.) can facilitate evolution of a single binding site.

Here, we focus on the 14 kDa cyanobacterial *Oscillatoria agardhii* agglutinin (OAA) lectin.^31^ OAA has a unique beta-barrel topology with two binding sites that each recognize the pentamannose M5 core present in all HMGs (Fig. 1A).^32–34^ This M5 core is the entire mannose content of the HMG, Man_5_GlcNAc_2_ (M5), while extensions of the D1, D2, and D3 arms create M6, M7, M8, and M9 (Fig. 1A, 1B).^35,36^ Because OAA broadly interacts with all HMGs, M5-M9, we hypothesized that OAA could serve as a scaffold for diverse variants with specificity for each HMG structure. OAA is a highly stable protein with a melting temperate of >80°C, reversible folding, and a glycan binding site composed of loops that are amenable to mutation without disrupting the protein’s fold.^31^ Prior studies of an OAA-family lectin have shown that alanine mutations to the loops in one lectin binding site can generate stable monovalent variants.^32^ As a bivalent lectin, OAA can be employed as an antiviral agent by targeting the N-glycans of viral coat proteins of SARS-CoV-2,^32^ HIV,^37,38^ and influenza,^33^ in addition to a glycobiology tool that captures tumor-derived extracellular vesicles bearing a high fraction of HMGs.^39^

To improve its prospect as a biotherapeutic or bioanalytical tool, we hypothesized that improving the affinity of OAA for specific glycans will both modulate its antiviral activity and facilitate its use as a sensitive molecular probe. Here, we aim to develop a lectin tool with novel binding capabilities, contribute to the understanding of lectin-glycan interactions,^40^ and establish techniques for reliable engineering of CBPs. Using the ligand bound structure as a starting point, we designed a large combinatorial mutation library for phage display against the M5 core. We designed the OAA library to focus on the mutation of 21 residues near the binding site of the M5 core while maintaining the primary binding contact between W77 and the central mannose (Fig. 1C, 1D). The residues are grouped into three sequence regions we denote as loop 1, 2, and 3. Because OAA is a bivalent lectin, we made three alanine mutations to the opposite binding site to remove its binding contribution and therefore only screen the interactions of site 1.^32^

In this study, we focus on the enrichment of phage displayed variants based on interaction with M5 (Fig. 1E). We enriched the library for four rounds then assessed the populations by amplicon sequencing for sequence enrichment and diversity. We isolated select variants of interest for binding assessment by biolayer interferometry (BLI) where we discovered a highly selective variant for M5. Combining a screen of point mutation variants with the crystal structure bound to the M5 core, we were able to identify a mechanism for how selectivity and higher affinity can be created in OAA and broadly in protein-glycan interactions. Upon restoring the bivalency of OAA with the discovered mutations, we generated highly improved HMG recognition tools. We further demonstrate the utility of our novel OAA variants as both tools in glycobiology and antiviral agents.

## Results

### Phage display of a combinatorial OAA library

To engineer a variant library capable of extensive binding site mutations we relied on a synthetically generated library. Here, we could modify OAA with defined amino acid ratios to maximize quality variants. The high structural integrity of the beta-barrel allowed us to confidently mutate much of the loop region, however, the possible variants far exceed the library size limitations of phage display. Therefore, we aimed for the naïve library to contain an average of 6 mutations per variant with a distribution between 1 and 13 mutations per variant (Fig. 2A, S1). We displayed OAA on M13 bacteriophage^41,42^ and conducted 4 rounds of enrichment against streptavidin beads bearing biotin-M5 with four bead washes as the only selection pressure. For each round of selection, we recorded the phage yield after binding in reference to a selection on beads without glycan. The yield peaked at round 2 with a 133-fold enrichment, decreasing to a 35-fold enrichment at round 4 (Fig. S2). Despite the yield plateau and drop, we proceeded with analysis of round 4 as it appeared to further obviate the enriched motifs seen in round 2 (Fig. S3). After the 4 rounds, 9 of the 21 mutated residues had at least 10% mutation adoption (Fig. 2A, S3). These sites are generally consistent with less conserved residues across the OAA lectin family (Fig. S4). However, residues Q31 and N75 are conserved through evolution, yet generated novel mutations in our phage library. The mutation-adopting residues are dispersed evenly throughout the three loops with exception of G55-G60 which also has strong resistance to mutation through evolution (Fig. 2B).

**Figure 2.**
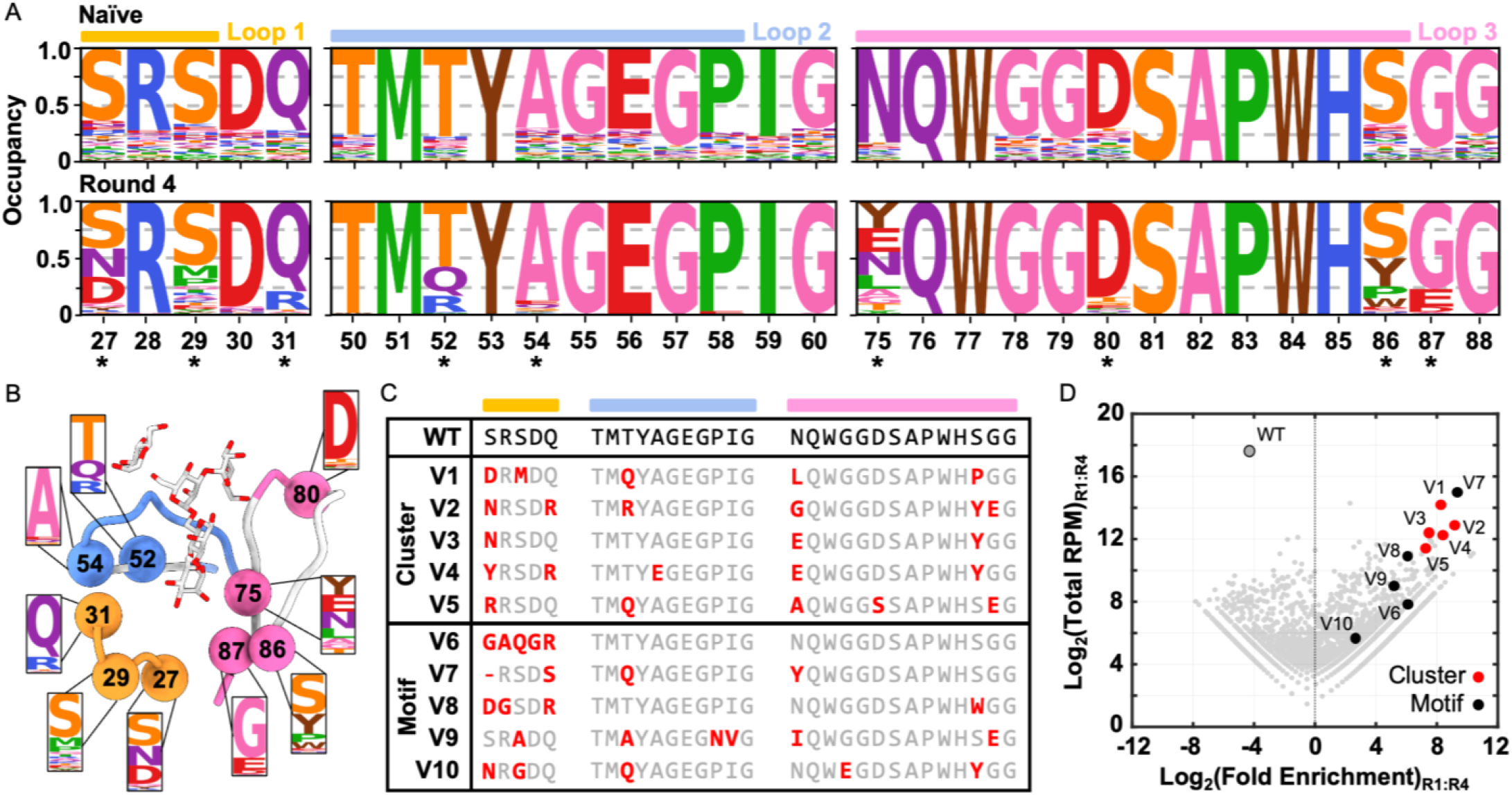
Enriched variant selection from phage display library. (A) (*top*) The sequence logo of the naïve library showing the distribution of amino acids at each mutation site. (*bottom*) The sequence logo of the library after round 4 of screening. Residues marked with an asterisk had at least 10% non-wildtype amino acids at the site. (B) A schematic of the asterisk residues located on the OAA binding site. (C) Sequence alignment of V1-V10 with wildtype OAA sequence. Mutated residues are marked in red. V1-V5 are grouped as the sequences derived from clustering. V6-V10 are grouped as the sequences derived from motif searches. (D) Volcano plot of the library changes from round 1 to round 4. The Log_2_(Total RPM) represents the total number of target reads per million of all reads in round 1 and round 4, while Log_2_(Fold Enrichment) represents enrichment from round 1 to round 4. Data along the top right of the plot will signify highly abundant and highly enriching variants. The cluster-based variants are marked in red, the motif-based variants are marked in black. The WT sequence is marked by a gray circle with a black outline.

The four rounds of enrichment reduced the library diversity 1000-fold, producing ∼2×10^5^ possible variants (Table S1). We chose to isolate probable high affinity variants from this library through amplicon sequence enrichment. Briefly, we clustered the sequences based on similarity (Fig. S5) to generate 10 clusters that represented 51% of the library reads. From these 10 clusters, we chose 5 for further assessment, V1-V5, ensuring that each enriched mutation site was present at least once (Fig. 2C). To access unique enriched sequences, we searched for small motif changes to find V6-V10 (Fig. S5). Plotting the abundance versus enrichment from round 1 to round 4 shows that V1-V5 all represent high abundance and enrichment variants, averaging a 250-fold enrichment. V6-V10 generally have lower abundance and lower enrichment with the exception of V7 potentially due to indel bias during amplification (Fig. 2D, S5).^43^ In contrast, the wildtype (WT) OAA sequence is detected to fallout 17-fold.

### Glycan binding screen of enriched variants

To evaluate the binding properties of each variant, V1-V10 were cloned into a pET28 vector, expressed in *E. coli*, and purified. We used bio-layer interference (BLI) to screen the binding of each variant to the following HMGs: M5, M6, M7D1 (M7), M8D1D3 (M8), and M9 (Fig. 3A, S6, S7). We observed a modest improvement in binding affinity (K_D_) for M5 in most variants, consistent with the phage enrichment against M5-coated beads. V4, V6, and V8 show the most notable binding affinity improvements. Generally, the variants decrease in binding affinity as sugars are added from M5 to M9. Some variants drastically reduce by the addition of a single sugar, for example, V2 and V4 from M5 to M6, V6 from M6 to M7, and WT from M8 to M9. These data indicate that glycan selectivity of M5 and M6 was generated during the selection process. From the kinetics standpoint, the highest affinity variant-glycan interactions appear to coincide with slower dissociation. For example, WT has a slow dissociation (k_off_) with M5-M7, V4 with M5, and both V6 and V8 with M5 and M6.

**Figure 3.**
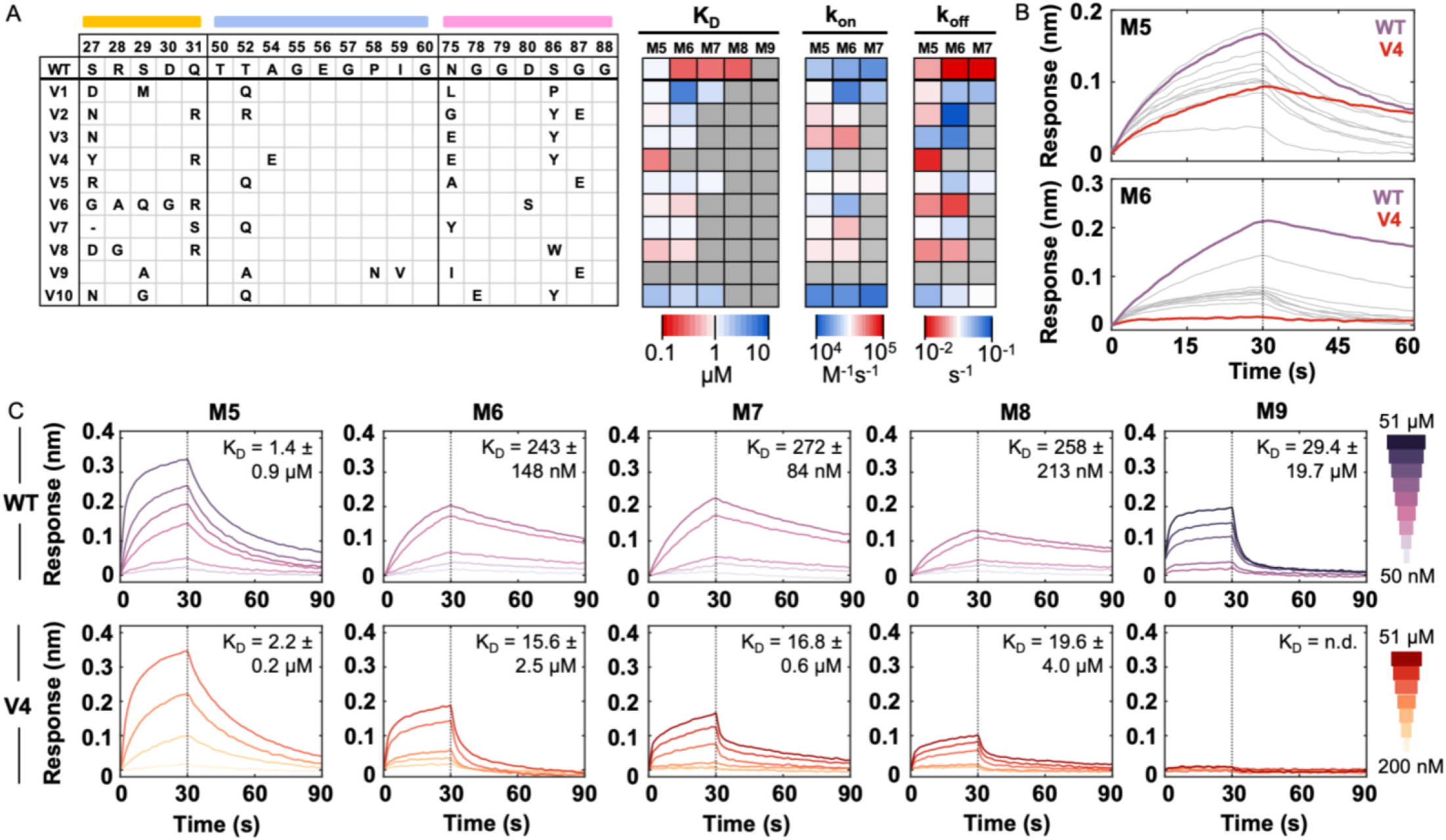
High mannose glycan interaction screen of selected variants. (A) Table of variant mutations with residue numbers and loop colors shown at top. Each variant was screened at 1 μM against each biotin-glycan, M5-M9. Heat maps of BLI data are shown for sensor response, binding affinity (K_D_), association rate (k_on_), and dissociation rate (k_off_). Heat map color red signifies better binding character, while blue signifies worse binding character. Gray boxes mark variant-glycan interactions that did not reach 0.05 nm binding response and therefore were not fit for kinetics. The K_D_ heat map shows a range of 0.1 (*red*) to 10 μM. The k_on_ heat map shows 1×10^4^ to 1×10^5^ (*red*) M^-1^s^-1^. The k_off_ heat map shows 1×10^-2^ (*red*) to 1×10^-1^ s^-1^. The values were determined by the average value of a duplicated BLI screen. (B) (*top)* BLI sensorgrams of WT OAA (purple) and V1-V10 (gray) against M5 plotting the sensor response over time. (*bottom*) BLI sensorgrams of WT OAA and V1-V10 against M6. V4 is highlighted in red. (C) (*top*) BLI sensorgrams for titrations of WT OAA with each glycan M5, M6, M7, M8, and M9, showing sensor response over time. Concentrations range from 50 nM to 51 μM. (*bottom*) BLI sesnsorgrams for titrations of V4 with each glycan M5-M9. Concentrations range from 200 nM to 51 μM. Inlaid in each plot are the binding affinities (K_D_), showing the average among duplicate titrations ± the standard deviation, where (n.d.) signifies not determined.

Comparison of all variant BLI results led to a standout variant, V4, which had slow dissociation and striking selectivity for M5 upon overlaying BLI sensorgrams (Fig. 3B). To elucidate these interactions beyond screening conditions, we completed binding titrations for both WT OAA and V4 (Fig. 3C, S8, Table S2). The K_D_ of V4 for M5 at 2.2 ± 0.2 μM remained within range of the WT K_D_ of 1.4 ± 0.9 μM. Therefore, the apparent dissociation rate improvement from WT (5.0 x 10^-2^ ± 1.7 x 10^-2^ s^-1^) to V4 (3.8 x 10^-2^ ± 7.8 x 10^-5^ s^-1^) was not significant enough to improve affinity for M5. On the other hand, compared to WT, V4 showed a remarkable 60-fold reduction in affinity for M6-M8 and no discernible interaction with M9. This translates to a 7-fold preference for M5 over M6, whereas WT has a 6-fold preference for M6 over M5. To determine the mutations responsible for selectivity we sought out structural evidence since the binding results did not reduce the complexity of mutations to a clear trend.

### Structural analysis of V4 bound to glycan

We determined a high-resolution crystal structure of the bivalent variant, V4V4 (PDB:10KB), bound to the M5 core, 3α,6α-mannopentaose (Fig. 4A, Table S3). We crystallized V4V4 rather than V4 to improve crystal contact symmetry and to compare mutation effects to both OAA binding sites. The structure shows that V4V4 binds the M5 core in the same orientation as previously determined for wildtype OAA, with an average Cα root mean squared displacement (RMSD) of 0.47 Å over the length of the protein (Fig. 4B). We also observe that sites 1 and site 2 show high similarity, with an overall Cα RMSD of 0.25 Å, suggesting that the properties of site 1 likely transfer to site 2 (Fig. S9). Regions of higher RMSD between V4V4 and WT are seen in the mutated loops. For example, in site 1 there is a RMSD of 2.0 Å for loop 1, 0.75 Å in loop 2, and 1.0 Å in loop Looking closer at binding site 1, immediate differences to the wildtype pocket are present in the sidechains of mutated residues, most strikingly in the orientation and hydrogen bond network of R28 (Fig. 4C). The N75E mutation extends the sidechain to form a salt bridge with R28, while the Q31R mutation releases the coordinating hydrogen bond that previously positioned both E56 and R28 towards the glycan. The A54E mutation does not appear to influence loop 2 or nearby sidechains, but rather the backbone carbonyl forms a new hydrogen bond with Q31R (Fig. 4D). Therefore, A54E and the nearby D30 may stabilize Q31R from interfering with the flipped R28 orientation. The R28 flip may only be possible because of the 2.0 Å inward shift of loop 1 (Fig. 4E) that is bolstered by the S27Y tyrosine, pointed opposite to the wildtype serine to accommodate the size (Fig. 4F). S27Y now packs onto the G87/G88 backbone and the S86Y tyrosine to restructure loop 1, previously shown through NMR structural restraints of WT OAA to have high conformational flexibility in the unbound state.^44^ Meanwhile, the R28 orientation may be important for shifting the D1 mannose 0.9 Å from the wildtype positioning (Fig. 4G). Together with the M6 mannose cavity being shrunken by S27Y, N75E, and R28, this new D1 arm position may diminish the ability of OAA to bind to the further extensions of M6 and M7 (Fig. 4H). Overall, the structure of V4V4 shows that OAA binding kinetics may be altered by the rotamer flip of R28 and the loop 1 shift, while selectivity is created by the shift and occlusion of the M6 binding site. Still, we aimed to elucidate the contribution of each mutation to understand how selectivity develops within a highly mutated carbohydrate binding site.

**Figure 4.**
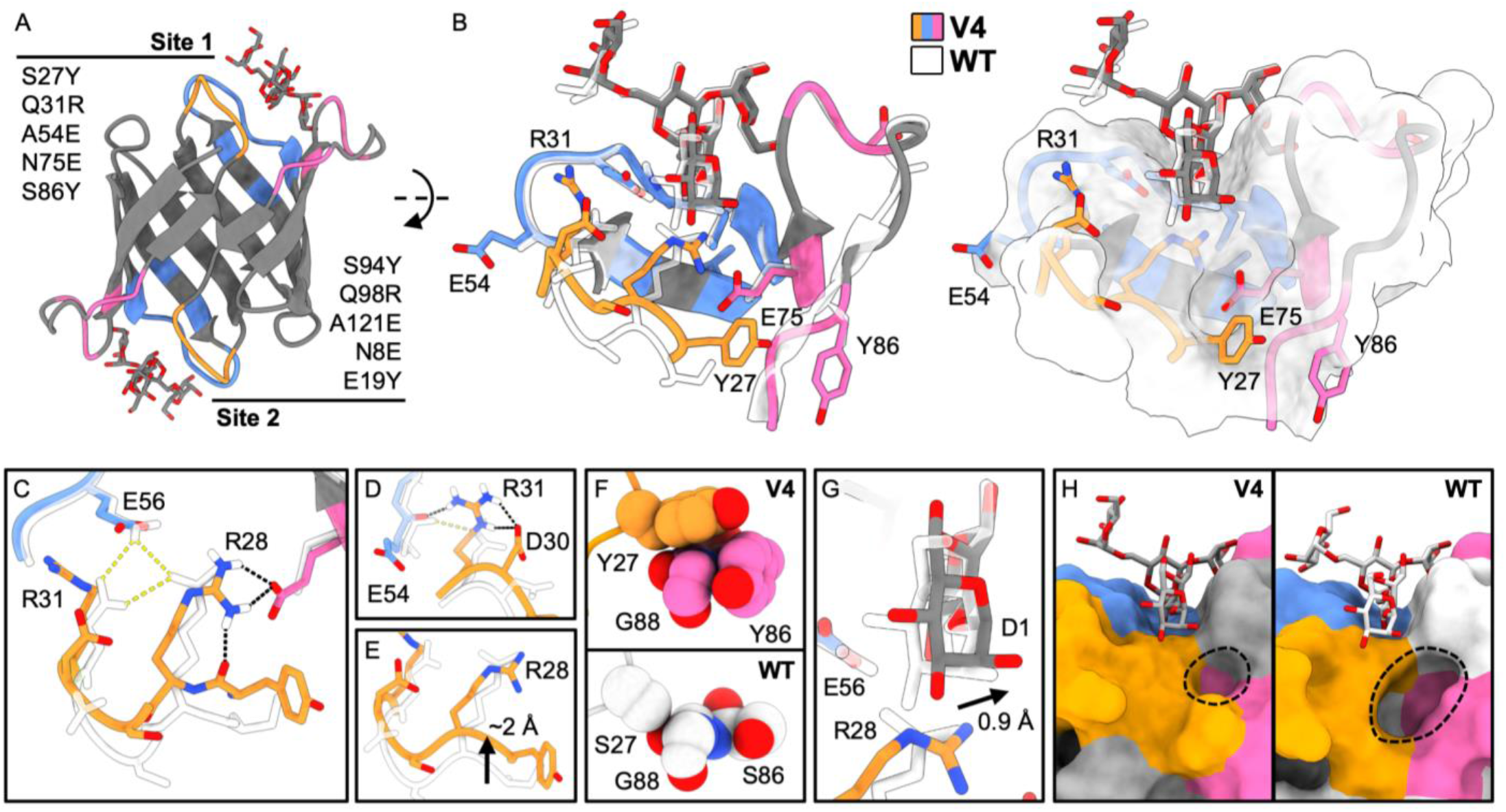
Structural effects of V4 mutations. (A) Beta-barrel structure of V4V4 (PDB:10KB) bound to 3α,6α-mannopentaose at site 1 and site 2 with loops colored by identity at both sites. Ligand is shown in gray with heteroatoms colored. Mutations made to each binding site are listed. (B) (*left)* Close-up of V4V4 site 1 with all library sidechains shown and colored by loop and heteroatom. Overlaid with WT OAA (PDB:3S5X) site 1 in white. (*right*) Overlay shown with WT OAA in surface representation. (C) Zoom-in of 4B showing the hydrogen bond network of loop 1 in V4V4 (black dashes) and WT OAA (yellow dashes). R28 makes new interactions with E75 and Y27 while releasing from Q31 and E56. (D) Hydrogen bonding of R31 which maintains the backbone interaction with E54. (E) Loop 1 undergoes a 2 Å shift inward from Y27 to S29. (F) (*top*) Zoom-in of Y27/Y86 packing onto loop 3 in V4 site 1. (*bottom*) Zoom-in of S27/S86 in WT OAA site 1. Atoms are displayed with Van der Waals radius spheres to accentuate packing. (G) The D1 arm of M5 is shifted 0.9 Å to the right, consistent with the R28 rotamer flip. (H) (*left*) Zoom-in of surface representation of V4 site 1. (*right*) Zoom-in of surface representation of WT OAA site 1. Potential binding pocket for the M6 mannose is circled.

### Mutational landscape of V4 Man-5 selectivity

To connect each mutation with functional consequence, we made 25 point-mutation (PM) variants containing between one and four of the mutations found in V4 (Fig. 5A, S10, S11). We screened PM1-PM25 against glycans M5-M9 using BLI to assess binding affinity and kinetics. Based on the trends in binding affinity, we binned the PMs into four groups: affinity enhancing (Fig. 5B), M5-selective (Fig. 5C), generally weakening (Fig. 5D), and destructive to all glycan interaction (Fig. 5E).

**Figure 5.**
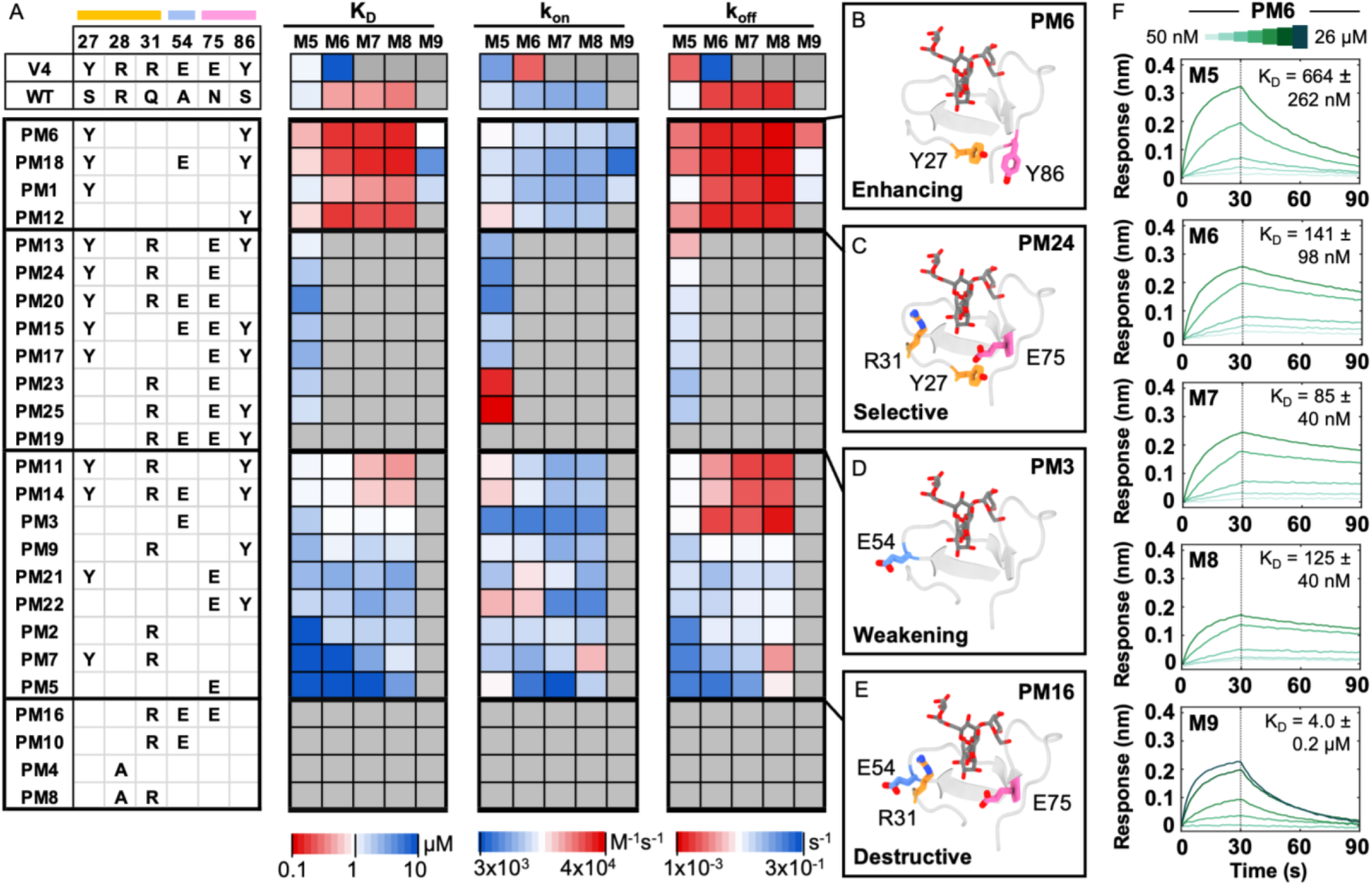
High mannose glycan interaction screen of V4 point mutation variants. (A) Table of V4-based point mutation variants with residue numbers and loop colors shown at top. Each variant was screened at 5 μM against each biotin-glycan, M5-M9. Heat maps of BLI data are shown for binding affinity (K_D_), association rate (k_on_), and dissociation rate (k_off_). Heat map color red signifies better binding character, while blue signifies worse binding character. Gray boxes mark variant-glycan interactions that did not reach 0.05 nm binding response and therefore were not fit for kinetics. The K_D_ heat map shows a range of 0.1 (*red*) to 10 μM. The k_on_ heat map shows 3×10^3^ to 4×10^4^ (*red*) M^-1^s^-1^. The k_off_ heat map shows 1×10^-3^ (*red*) to 3×10^-1^ s^-1^. The values were determined by the average value of a duplicated BLI screen. (B-E) The following are mutations mapped onto the V4V4 structure for representative variants of each binned group. (B) Mutation positions for enhancing variant, PM6. (C) Mutation positions for selective variant, PM24. (D) Mutation positions for weakening variant, PM3. (E) Mutation positions for destructive variant, PM16. (F) BLI sensorgrams for titrations of PM6 with each biotin-glycan M5, M6, M7, M8, and M9, showing sensor response over time. Concentrations range from 50 nM to 26 μM. Inlaid in each plot are the binding affinities (K_D_), showing the average among duplicate titrations ± the standard deviation.

The enhancing group exhibited higher binding affinity across all glycans. At least one of the tyrosine mutations are present in each enhancing variant, the greatest impact seen when both are present. PM6 (S27Y, S86Y), which consists of the two tyrosine mutations, showed a remarkable affinity enhancement for M9. We performed full binding titrations with PM6 to confirm an ~8-fold lower K_D_ for M9 and a K_D_ below 100 nM for M7 (Fig.5F, Table S2). PM1 (S27Y) shows that the S27Y mutation alone confers improved binding to M9, suggesting the loop 1 shift as the driving factor. We also tested (S27F, S86F) and (S27W, S86W) variants to find that any aromatic side chain is sufficient for affinity enhancement and therefore the loop 1 shift (Supplemental Data 2).

The tyrosine mutations are also prevalent in the selective and weakening groups. In PM17 (S27Y, N75E, S86Y), the addition of N75E to the tyrosines converts the site to M5 selectivity. By also adding Q31R, the selectivity is even further improved by decreasing the affinity for M6-M9 as in PM13 (S27Y, Q31R, N75E, S86Y). Altogether, S27Y appears most influential because PM24 (S27Y, Q31R, N75E) is the minimal unit that confers similar binding properties to V4. This agrees with the proposed function of S27Y to position loop 1 for R28 to form a salt bridge with N75E and shrink the M6 pocket.

The weakening group is generally composed of single mutations or combined mutations that are incapable of M5 selectivity. When the mutations N75E and Q31R are alone they move into the weakening group, as seen in PM2 (Q31R) and PM5 (N75E). Interestingly, the N75E which we posit as responsible for the R28 rotamer flip, shows no selectivity on its own, emphasizing the requirement of Q31R in PM23 (Q31R, N75E) to induce selectivity. The destructive variants show the importance of the tyrosines for providing binding capability. For example, PM16 (Q31R, A54E, N75E) which represents V4 without S27Y and S86Y had low expression yield and had no detectable binding. These data suggest that the mutations responsible for remodeling the hydrogen bonding network of the binding pocket are deleterious on their own and require co-dependent mutations for their effect.

The k_on_ and k_off_ can also provide insight into the mutation sites. Broadly, the dissociation rates appear to dictate the overall binding affinity. For example, the entire enhancing group has slower dissociation rates than WT but varies in association rate. PM12 (S86Y) shows that S86Y offers improved kinetics, potentially through stabilizing loop 3 which positions the central W77. While PM1 (S27Y) shows that S27Y has little effect on kinetics in M5-M8 but allows improved binding to M9. The selective group generally has worse k_on_ and k_off_ values except for PM13 (S27Y, Q31R, N75E, S86Y) which most closely resembles V4. The weakening group supports the prior assessment of S86Y improved association rates and shows contributions of single point mutations. For example, PM3 (A54E) shows a slow association rate consistent across M5-M8, while PM2 (Q31R) shows an above average association rate across M5-M8.

### Transferability of variant mutations onto site 2

We have generated two variants of OAA with especially useful properties, V4 with selectivity for M5, and PM6 with improved affinity for all HMGs. We determined the site 1 binding affinity for M5-M9 to identify these variants. To convert these variants into bivalent tools, the site 2 mutations need to exert the same function. An overall comparison of the two binding sites shows high similarity (Fig. 6A, S9), which we verified by screening the interactions of site 2 monovalent variants (Fig. S12). We therefore wondered how the binding of the second site would influence the affinity enhancement or glycan specificity. When both sites contain PM6 mutations, PM6PM6 has a binding affinity for M9 of 22 ± 3 nM, ∼26-fold stronger than WTWT (Fig. 6B, S13). For reference, the affinity of PM6 for M9 is ∼8-fold greater than WT. This exhibits that the tyrosine mutations also improve site 2 despite not having complete site conservation (Fig. S14). Shockingly, the V4V4 mutant exhibited a drastic selectivity improvement over the V4 mutant. Affinities for M5, M6, and M7 were 18.04 ± 0.01 nM, 3.69 ± 0.04 μM, and 13.07 ± 0.34 μM respectively, where the error is the fitting uncertainty derived from a single titration (Fig. 6C, S13). While the V4 mutant has a 7-fold preference for M5 over M6, V4V4 shows a remarkable 205-fold preference of M5 over M6. Impressively, there is still no detectable binding to M9 despite the avidity-based enhancement of bivalency.

**Figure 6.**
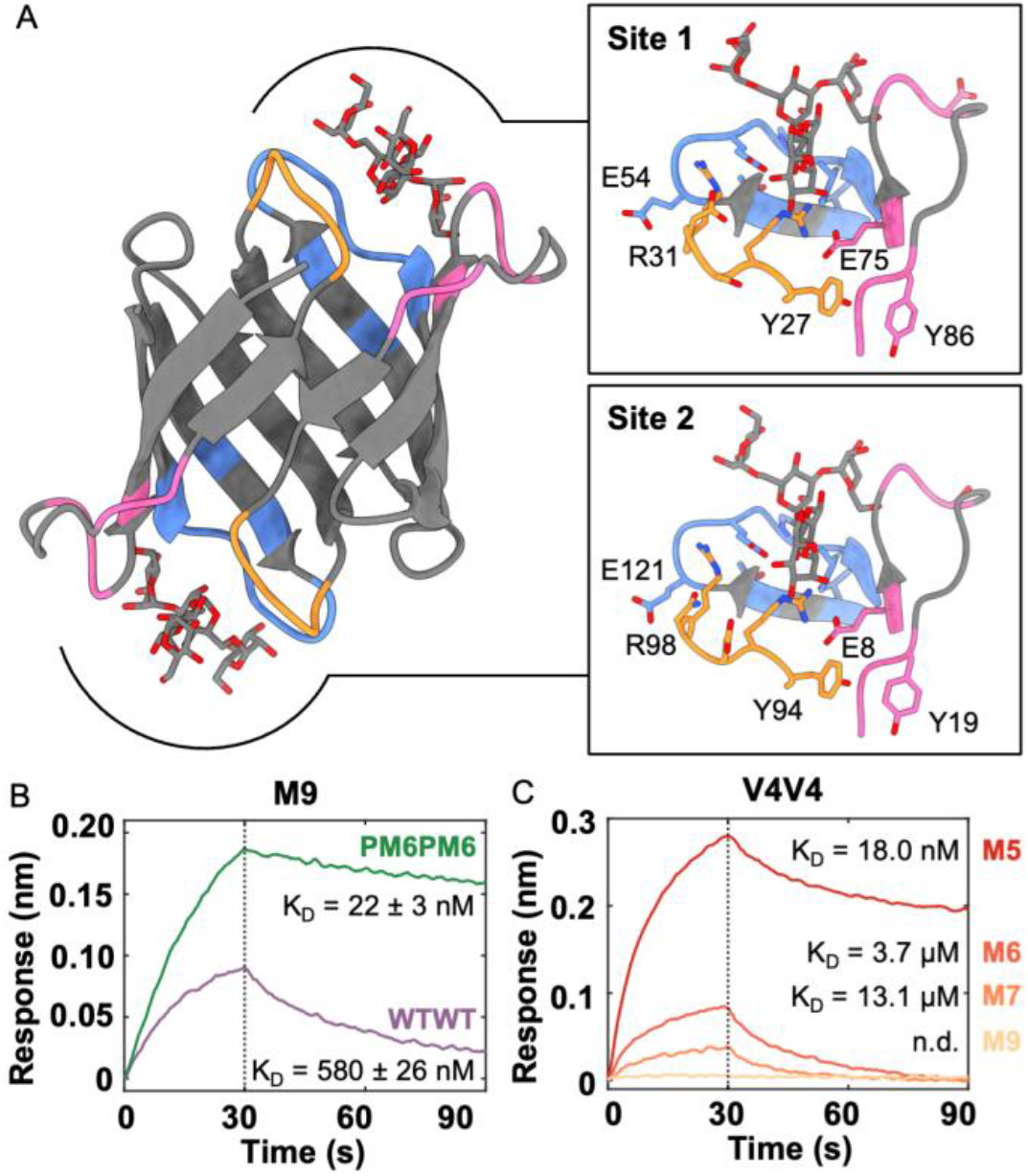
Amplification of variant functionality by bivalency. (A) (*left)* Beta-barrel structure of V4V4 (PDB:10KB) bound to 3α,6α-mannopentaose at site 1 and site 2 with loops colored by identity at both sites. (*right*) Close-up of site 1 (*top*) and site 2 (*bottom*). (B) BLI sensorgrams of 1 μM WTWT (purple) and 1 μM PM6PM6 (green) against M9 plotting the sensor response over time. Inlaid are the binding affinities (K_D_) ± the standard deviation derived from duplicate titrations. (C) BLI sensorgrams of 1 μM V4V4 against M5 (*red*), M6 (*light red*), M7 (*orange*), and M9 (*light orange*) plotting the sensor response over time. Inlaid are the binding affinities (K_D_), where (n.d.) signifies not determined.

### Applications of affinity and selectivity enhanced OAA variants

Finally, we sought to demonstrate the utility of our engineered variants in biological systems as glycoform separation tools and viral inhibitors. We used RNase B as a model system to test how our variants could capture specific protein glycoforms. RNase B has a single N-glycosylation site with only the M5-M9 HMGs. We started by characterizing the affinity of each monovalent variant for RNase B, which is predominantly M5 glycosylated.^45^ We determined the binding affinities to be 137 ± 22 nM for WT, 1020 ± 11 nM for V4, and 78.5 ± 1.3 nM for PM6 (Fig. 7A, S15). These are consistent with the affinity range we determined for the biotinylated glycans. V4 has a markedly lower affinity for RNase B due to the weak interaction with the M6-M9 present. After confirming our variants translate to glycoprotein interactions, we conducted pulldown assays followed by intact mass spectrometry to identify the RNase B glycoforms bound by WT, V4, and PM6. We characterized the starting RNase B material to contain roughly 45% M5, 25% M6, 5% M7, 20% M8, and 5% M9 (Fig. 7B, Table S4). In this case, we know the number of sugars per glycan but cannot differentiate the exact linkages for M7 or M8. For example, while our biotinylated glycan assays only use M7D1, both M7D2 and M7D3 are also present in RNase B. The pulldown results for WT show a slight increase in the M6-M8 fractions compared to input, and a strong loss of M9 (0.3% in pulldown versus 5% in input). V4 binds predominantly M5 (96.7% of pulldown) with minor M6 and M8 peaks. PM6 shows a similar profile to WT, except for a 5-fold increase in the M9 peak at 1.5% abundance. The pulldown results match the affinity trends of the BLI data and illustrate the striking ability of V4 to separate only M5 glycoforms from a pool of high mannose glycans.

**Figure 7.**
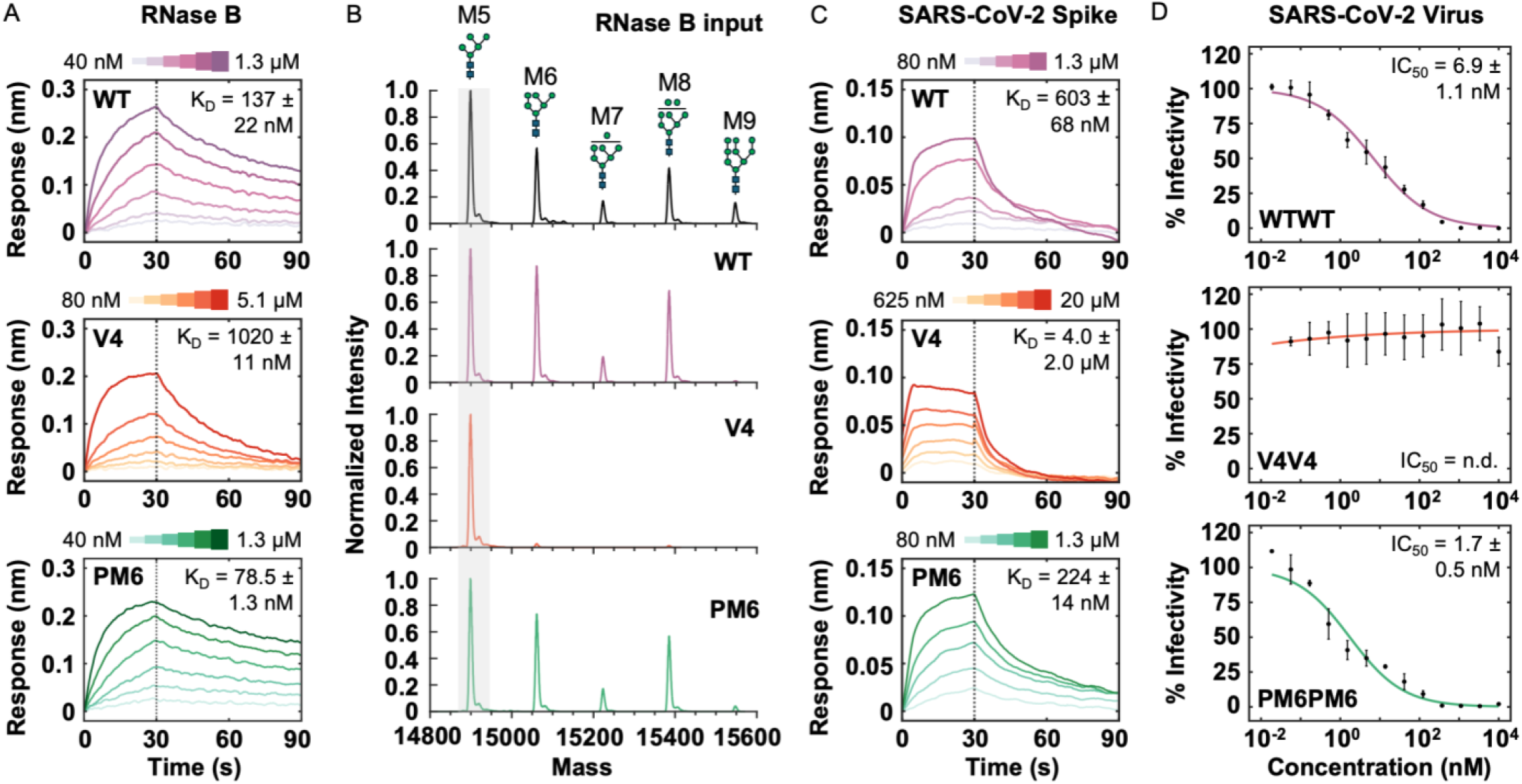
OAA variants as improved glycan-targeting tools. (A) BLI sensorgrams of titrations for His-tagged WT OAA, V4, and PM6 against RNase B showing sensor response over time. Concentrations range from 40 nM to 1.3 μM (WT), 80 nM to 5.1 μM (V4), and 40 nM to 1.3 μM (PM6). Inlaid in each plot are the binding affinities (K_D_), showing the average among duplicate titrations ± the standard deviation. (B) Mass spectra for RNase B input and the pulldowns of WT OAA, V4, and PM6. Peaks are present for M5, M6, M7, M8, and M9 with possible structures shown above. Peaks are normalized to a M5 peak intensity of 1.0. (C) BLI sensorgrams of titrations for His-tagged SARS-CoV-2 Spike against WT OAA, V4, and PM6 showing sensor response over time. Concentrations range from 80 nM to 1.3 μM (WT), 625 nM to 20 μM (V4), and 40 nM to 1.3 μM (PM6). Inlaid in each plot are the binding affinities (K_D_), showing the average among duplicate titrations ± the standard deviation. (D) Concentration-dependent inhibition of SARS-CoV-2 by OAA variants, determined by a focus reduction neutralization assay. Concentrations are plotted using a log_10_ scale and the normalized foci counts are reported as % infectivity of virus. Biological triplicate measurements were fit to a dose-response curve to achieve IC_50_ values for each variant, where (n.d.) signifies not determined.

OAA family lectins were first characterized for their antiviral properties towards viruses such as influenza, HIV and SARS-CoV-2.^32,34,46,47^ We began by determining the affinities of the monovalent variants for the SARS-CoV-2 spike glycoprotein, which would be the target of OAA during viral inhibition. We determined a K_D_ of 603 ± 68 nM for WT, 4.0 ± 2.0 μM for V4, and 224 ± 14 nM for PM6 (Fig. 7C, S15). The SARS-CoV-2 spike has immense glycan heterogeneity among its 22 conserved sites of N-glycosylation which are predominately occupied by complex and hybrid glycans.^48,49^ There are five sites occupied by high mannose glycans,^50^ of which are largely M5/M6/M7 glycans, and an additional three sites displaying either hybrid glycans or a M5 glycan.^51^ We sought to determine if the affinity differences between the OAA variants could modulate viral entry inhibition. For these assays we used the dimeric variants because monovalent OAA is incapable of the multivalent-driven agglutination mechanism.^32,52–54^ We observed potent inhibition for both WTWT and PM6PM6, with an IC_50_ of 6.9 ± 1.1 nM and 1.7 ± 0.5 nM, respectively (Fig. 7D, S16). In contrast, V4V4 showed no inhibition within the concentration range of 56 pM to 10 μM. To benchmark the efficacy of our OAA variants, we compared them to a previously characterized OAA-family SARS-CoV-2 inhibitor, BOA. BOA has four glycan binding sites instead of two, leading to a highly effective IC_50_ of 0.136 ± 0.003 nM (Fig.S16).^32^ While BOA showed a 51-fold better inhibition than WTWT, we were encouraged to find that BOA was only 12-fold better than PM6PM6, a lectin with half the size and valency. We conclude that engineering the OAA lectin for higher affinity and unbiased HMG interaction is a promising strategy for viral inhibition and detection,^55^ while selectivity engineering for these purposes will require the viral glycoprotein to be predominantly modified by the target glycan.

## Discussion

The engineering of CBPs has been limited by the challenging requirements for glycan specificity.^24^ Our study supports a path for phage display to tune lectins toward selective glycan binding.^56^ We achieved two distinct selective variants, V4 and V6, without requiring external selection pressure to acquire selectivity (Fig. 8, Table S2). We focused on V4, a 5-mutation variant that we dissected to reveal how co-dependent mutations confer M5 selectivity. By isolating the enhancing mutations, we generated a new variant, PM6, which had a marked improvement in affinity for M9 and overall interaction with HMGs.

**Figure 8.**
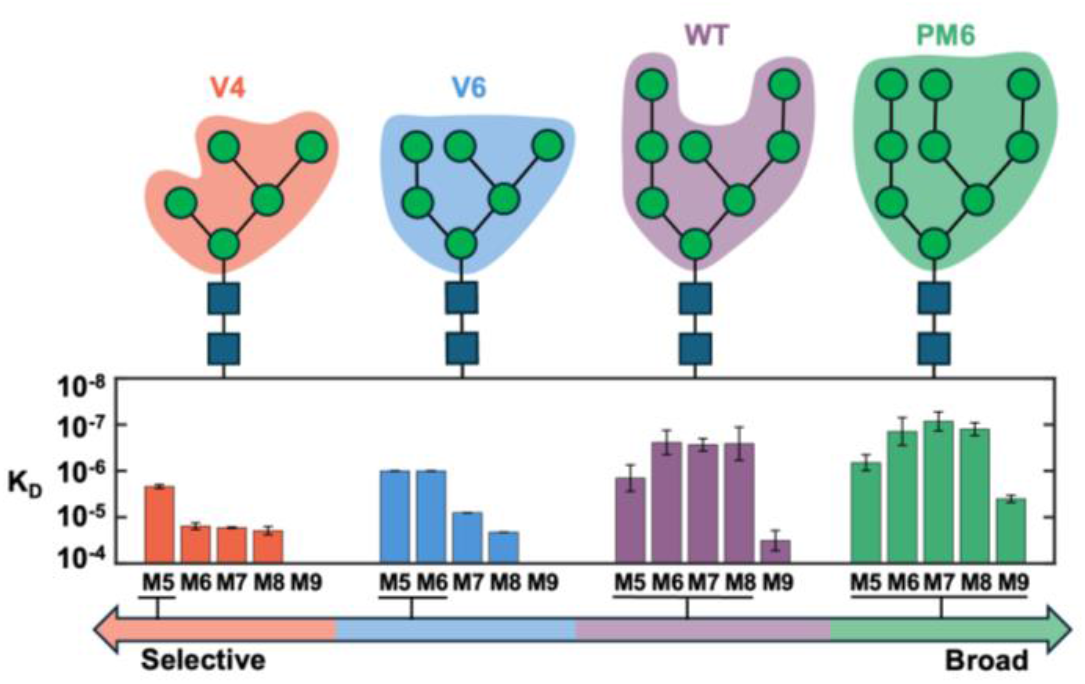
HMG preferences of engineered OAA variants. Binding affinity expressed as K_D_ (M^-1^) for each variant against M5-M9, plotted with a log_10_ scale. Variants V4 and V6 discovered from phage display enrichment yielded new HMG selectivity for OAA, while the point mutation dissection of V4 yielded a broadly interacting high-affinity variant, PM6. Binding preferences indicate these variants have low micromolar affinity or better with only the designated glycans.

The success of our library was likely achieved because OAA is a stable beta-barrel allowing for a high number of mutations to the loop regions. V6 for example, acquired a complete loop 1 swap and maintained wildtype-level affinity. OAA also proved stable for the site 2 knock-out used for phage display, enabling a monovalent OAA lectin necessary for disentangling site 2 effects. Amazingly, the conserved symmetry of OAA allowed us to revert to bivalency by transferring site 1 mutations to site 2. This strategy not only allowed for single-site optimization but also the manipulation of avidity effects, which many lectins use to overcome their weak single-site affinity for carbohydrates in the μM to mM range.^57^ Our library design allowed us to push the limits of a single carbohydrate recognition domain, reaching a K_D_ as low as 85 nM for M7, then reintroduce the native valency. In PM6PM6 for example, we observed a low-nanomolar binding affinity for its least preferred HMG, M9, a 26-fold enhancement from the wildtype. This places PM6PM6 as a potential high affinity candidate for unbiased interaction with the entire class of HMGs. On the side of selectivity, reverting to bivalency also amplified the effects of V4. Conversion to V4V4 took the 7-fold preference for M5 over M6 to an outstanding 205-fold preference, a 727-fold preference for M5 over M7, and altogether remained incapable of binding M9. This level of glycan specificity transitions lectins into precision tools for an entire glycan structure rather than just sub-glycan motifs or broad glycan types. Additionally, because we show site 1 and site 2 are independent, we look forward to future constructs with asymmetric binding preferences to link distinct targets.

Looking deeper at the variants extracted from phage display, V2 and V4 share M5 selectivity and the mutations S27N/Y, Q31R, N75G/E, and S86Y. However, V3 has the S27N, N75E and S86Y mutations with no M5 selectivity. Based on our mutational screen, we propose this is due to the lack of Q31R and the tyrosine of S27Y. In the case of V2 which lacks N75E, we propose that selectivity is acquired by its unique G87E mutation that points directly toward the N75G site (Fig. S17). This transfers the selective glutamate from N75E to the G87 site and satisfies the R28 rotamer flip from a different angle. A similar story appears with Q31R, as seen in V6 and V8 which are selective for M5 and M6. These variants differ from V2 and V4 by replacing R28 with the Q31R mutation to maintain glycan interaction, thereby relocating the essential arginine to another site on loop 1. This was verified by PM8 (R28A, Q31R) which showed that Q31R alone could partially rescue the R28A mutant for M5 and M6 binding. Together, we show that selectivity is generated through mutation of two highly conserved residues Q31 and N75, and one evolutionarily invariant residue, R28 (Fig. S4). Our combinatorial mutation library circumvents stepwise evolutionary pathways that would not persist through the binding-deficient mutation of N75E necessary for selectivity. We take away a powerful engineering tactic that loss of function mutations can be recovered by co-mutations of nearby residues, therefore conferring new binding affinities, kinetics and selectivity.

We were able to explore another engineering strategy through the dissection of V4 to discover PM6, a variant exhibiting higher affinity for a broadened range of HMGs. Particularly, PM6 had low micromolar affinity for M9 which includes the terminal D2 sugar. This sugar regularly precludes strong WT OAA interaction to M7D2, M8D1D2, M8D2D3, and M9. Interestingly, although we hypothesized that our high affinity variants would either add hydrogen bonding contacts via polar side chains or ring stacking via new aromatic residues, PM6 instead enhanced affinity by shifting loop 1 inward. Therefore, rather than changing the direct contact residues, mutations distal to the ligand can buttress loop positioning to improve the wildtype-ligand interaction network. A distal tyrosine mutation was also found to improve the affinity of a different lectin CV-N, suggesting this may be a design goal for stabilized CBP binding sites.^58^ We would like to note that these positive effects can be offset by weakening mutations to generate selective binding that would otherwise be incapable of ligand binding. As a result, we imagine V4 as an unlikely product of rational design, computational design or even directed evolution due to the mutational pathway required toward selectivity.

Engineering of CBPs has generally oriented toward tool development for glycomics.^59,60^ V4 can be immediately useful for this application as a M5-specific tool. As shown by the RNase B pulldown, V4 enriches samples for M5-bearing glycoproteins. M5 is a lingering mark on glycoproteins in serum, as M6-M9 glycans are enzymatically reduced to M5 by serum mannosidases.^61^ M5 can impact the pharmacokinetics of antibodies, both marking it for clearance while also improving protein stability and antigen interaction.^62–64^ On the other hand, PM6 can be used to extract all HMGs with little bias, adding a new tool to quantify high mannose content. High mannose content is typically associated with immature processing within the Golgi, they are therefore a biomarker for cases of abnormal protein production like in cancers and viral replication.^65^ Importantly, because OAA utilizes the M5 core there should be little off-target binding to complex or hybrid glycans containing mannose. We therefore envision PM6 as a new tool to track cancer biomarkers,^17^ detect viral infections,^66^ and identify antibody glycosylation.^67^

We also imagine OAA variants as attractive candidates to take on the role of glycan-targeted antiviral therapeutics. Glycans typically lack immunogenicity, making them a difficult target antigen to generate antibodies.^68,69^ Antiviral CBP engineering has focused on a small of set of well-characterized lectins like CV-N^58^ and BanLec,^70–72^ in addition to some glycan-targeting antibodies like 2G12.^73–75^ Here, we showed that our affinity enhanced bivalent variant PM6PM6 translated into a 4-fold improved viral inhibitor. The improvement of PM6PM6 over WTWT demonstrates the coupling between affinity enhancement and viral inhibition. By nearing the inhibitor potency of BOA, a lectin with double the valency of OAA, we demonstrated that sequence space facilitates higher binding affinities that may not be selected for by natural selection of CBPs which often rely on high valency. Meanwhile our M5-selective variant, V4V4, which we showed to have an 18 nM binding affinity for M5 was unable to inhibit SARS-CoV-2 at concentrations up to 10 μM. This stems from the proposed mechanism of OAA-mediated viral inhibition, where a single OAA protein links together two separate viral spike proteins.^32,76^ Because of the glycan heterogeneity deposited onto the SARS-CoV-2 spike protein, it may be unlikely for V4V4 to locate a nearby spike protein with a M5 modification at the correct site. This emphasizes the importance of broad glycan interaction for efficient viral inhibition and the requirement for high abundance of specific glycans for selective lectins like V4V4 to be effective.

## Conclusion

We demonstrated that monovalent OAA is a powerful scaffold for generating HMG-specific CBPs. Our combinatorial library strategy effectively captures an expanded sequence space of OAA to produce novel phenotypes not selected for by nature. These unique variants can be transformed into highly effective bivalent tools for glycoform separation or viral inhibition. We hope that continued engineering efforts continue to use the underexplored scaffolds of lectins and glycotransferases to facilitate designer CBPs that rival the expansive diversity of glycans.^77–79^

## Acknowledgements

The authors would like to acknowledge and thank Dr. Tatiana Mishanina for help with plasmid electroporation, Dr. Yongxuan Su at the Molecular Mass Spec Facility (MMSF) at UCSD for data collection, and Dr. Jake Bailey at UCSD Crystallography facility for help with initial crystal screening.

Crystallization at the National Crystallization Center at HWI was supported through NIH grant R24GM141256. This work is based upon research conducted at the Northeastern Collaborative Access Team beamlines, which are funded by the National Institute of General Medical Sciences from the National Institutes of Health (P30 GM124165). The Eiger 16M detector on the 24-ID-E beam line is funded by a NIH-ORIP HEI grant (S10OD021527). This research used beamtime awards (DOI: https://doi.org/10.46936/APS-191148/60014907) from the Advanced Photon Source, a U.S. Department of Energy (DOE) Office of Science User Facility operated for the DOE Office of Science by Argonne National Laboratory under Contract No. DE-AC02-06CH11357. A.J.G. is supported by NIH grants R00GM145970 and U54CA272220. K.D.C. is supported by NIH grant R35GM144121. E.H. is supported by T32GM139795. VTM acknowledges support from the HHMI Gilliam Postdoctoral Fellowship The following reagent was obtained through BEI Resources, NIAID, NIH: SARS-Related Coronavirus 2, Isolate New York-PV08410/2020, NR-53514. The authors declare no conflicts of interest.

