## Supplemental Information for "Engineered OAA lectins as selective and sensitive high mannose glycan targeting tools"

**Materials and Methods**

**Constructs**

The sequence used for WT OAA (monovalent) is:

SGGGSALYNVENQAGGSSAPWNEGG**QWEIGSRSDQNVVAINVESGDDGQTLNGTMTYAGEGPIGFRATLLGNNSYEVENQWGGDSAPWHSGGNWIL**GSAENQNVVAINVESGDDGQTLNGTMTYAGAGPIGFKGTLT

Where a SGGGS N-terminal linker was used in both pComb3XSS and pET28b for all variants and constructs. In bold is the amplicon range used for sequencing. A full detail of all variant sequences used in this study can be found in Supplemental Data 1.

**Library Generation and Cloning**

To display the OAA variants, we used the phagemid pComb3XSS (pComb3XSS was a gift from Carlos Barbas (Addgene plasmid #63890 ; http://n2t.net/addgene:63890 ; RRID:Addgene_63890).^1^ The OAA combinatorial mutagenesis library insert was synthesized, cloned into pComb3XSS, and sequenced by GenScript (New Jersey, USA and Zhenjiang, China). The domain structure during display is OmpA signal peptide-OAA-6xHis-HA-pIII, where OmpA locates the construct to the periplasm and pIII is a coat protein of M13 bacteriophage. The phagemid library was electroporated into XL1-Blue competent cells (Agilent) using a Eporator (Eppendorf) with 1 mm/90 μl electroporation cuvettes (Fisher). Briefly, XL1-Blue cells were grown in 2xYT (Fisher) + tetracycline (Fisher) until OD600 = 0.5. Cells were pelleted and washed twice with cold MQ H_2_O. Finally, the pellet was resuspended to an OD600 of 100, aliquoted to 50 μl and kept on ice until electroporation. 5 μl of 10 ng/μl of the library was added to the cell aliquot and electroporated at 1800 V/5 ms pulse. Samples were recovered in 950 μl of SOC media (Fisher) and shaken at 37 °C for 60 minutes. At this point, a 20 μl aliquot was removed to count colony forming units (CFU), and the rest was added to 10 times the volume of 2YT + carbenicillin + 2% glucose and shaken for 16 hours at 37 °C. The cells were spun down at 4,000xg and resuspended in 40% glycerol and stored at -80 °C to be used later for phage production.

**Phage Expression and Purification**

Phage production and purification were carried out as described by prior published protocols using M13 bacteriophage.^2–5^ Phagemid-containing XL1-Blue cells were inoculated in 500 mL 2xYT + 100 μg/mL carbenicillin/10 μg/mL tetracycline + 2% glucose to a starting OD_600_ below 0.2. The quantity of cells inoculated was chosen to give 100x the library diversity. The flask was shaken at 37 °C for several hours until OD_600_ = 0.8. Then, add 5x the number of cells (using OD_600_ = 1 = 8 x 10^8^ cells/mL as an estimate) of M13KO7 helper phage. Flasks were incubated for 25 minutes at 37 °C without shaking, followed by 15 minutes of shaking. Cells were spun down at 4,000 x g for 10 minutes and resuspended in 500 mL of 2xYT + carbenicillin. The flask was shaken for 2 more hours at 37 °C until 0.1 M IPTG was added alongside kanamycin to a final concentration of 25 μg/mL. The growth was shaken for 16 hours at 30 °C. After phage growth, the cells were spun down at 4,000 x g for 20 minutes at 4 °C. To purify the phages, the supernatant was recovered and 1/4^th^ of the volume of 5x PEG/NaCl (2.5 M NaCl, 20% w/v PEG8000) was added and left on ice for 1 hour. The mixture was then centrifuged at 12,000 x g for 1 hour, the pellet was resuspended in 50 mL 1x PBS. The resuspension was spun down at 4,000 x g for 20 minutes to remove any remaining bacteria. 1/4^th^ of the volume of 5x PEG/NaCl was added to the supernatant and left on ice for 30 minutes. Then the sample was spun down at 12,000 x g for 30 minutes and the pellet was resuspended in 1x PBS to roughly 10^12^ – 10^13^ CFU/mL as estimated by A269 absorbance. The phage stock was filtered through a 0.45-micron filter, and a phage titer assay was done to verify the CFU/mL. The phage assay was done on both carbenicillin and kanamycin plates to confirm that the phage sample had over a 10:1 ratio of OAA-phage:M13KO7 phage. After determining the phage concentration, glycerol was added to 40%, and phages were stored at -80 °C for long term storage or used immediately for the next round of selection.

To count the phage titer, we adapted the following protocol for spot plating.^4^ Briefly, a single colony of XL1-Blue cells was grown in LB + tetracycline until OD600 = 0.6 – 1.0. The cells could then be stored on ice until further use. In a 96-well plate serial dilutions were made of the produced phages in PBS. 10 μl of the desired dilutions (10^3^ – 10^8^) were combined with 90 μl of XL1-Blue cells. Phages and XL1-Blue cells were incubated for 15 min at 37 °C to allow the phage to attach to the cells. 5 μl of each dilution was triple spotted on 2YT agar plates supplemented with carbenicillin (phagemid) or/and kanamycin (M13KO7) and incubated overnight at 37 °C. Next day, the colonies were counted and calculated for overall CFU/mL.

**Phage Display on Magnetic Beads**

Selection and enrichment of the phage library followed the prior published protocols.^6,7^ Starting from a phage stock prepared from the naïve library or prior round of selection, 10^11^ CFU of phages were diluted to 1 mL in 1x PBS + 3% BSA. This number of phages was at least 100x the diversity of the library. To block background binding of phage to the tubes and Dynabeads™ M-280 Streptavadin (Invitrogen), all materials were pre-blocked with 1x PBS + 3% BSA + 0.1% Tween-20. All following steps with beads were performed at 25 °C. The phage library was incubated in a BSA-blocked tube for 30 minutes, then the library was incubated with pre-blocked ligand free beads for 30 minutes. The supernatant was removed and incubated with pre-blocked beads loaded with biotin-M5 (Chemitope Glycopeptide) for 1 hour. The supernatant was discarded and the beads were washed four times. Each wash step took roughly 1 minute, where beads are resuspended in 1x PBS, placed into a new pre-blocked tube, and placed on the magnetic rack for removal of supernatant. After the final wash, the beads were incubated in 500 μl of 0.1 M glycine-HCl pH 2.2 for 10 minutes to release bound phage. Then the supernatant was removed and placed into the final tube which contained 500 μl of 1x PBS + 0.2 M Tris-HCl pH 8.0. The phage titer was determined as previously described. Phage yield was quantified using a ligand free bead as a negative control and used as a reference population.

The phage eluate was stored at 4 °C until it was used to infect XL1-Blue cells, ensuring there a ratio over 10:1 cells:phage. The entire eluate was added to 2-20 mL of XL1-Blue cells (at OD600 = 0.6), depending on phage titer. This was then incubated at 37 °C for 25 minutes, then shaken at 37 °C for 15 minutes. The cells were spun down at 4,000 x g for 10 minutes. The supernatant was removed and the pellet was resuspended in 2 mL 2xYT. The resuspension was plated on a 245 x 245mm Nunc™ Square BioAssay Dish containing 2xYT + agar + carbenicillin + 1% glucose. The plate was left to grow colonies for 16 hours at 37 °C. After growth completion, 5 mL of 2xYT was added to the plate and the bacteria film was gently scraped off. The OD_600_ of the cell solution was taken to determine the cell count to determine the volume to be used for the next round of phage production to ensure there are at least 100x the number of phage diversity. At this point, the infected cells could be used directly for phage expression or frozen at -80 °C in 15% glycerol.

**Amplicon Sequencing and Analysis**

Plasmids were purified using Monarch Plasmid Miniprep Kit (New England Biolabs) from XL1-Blue cells infected by the phage eluate. The amplicons were PCR amplified using Q5 High Fidelity DNA Polymerase (New England Biolabs) with primers SeqF:GAAGGTGGTCAGTGGGAGATC (forward) and SeqR:CCGCAGAACCCAGAATCCA (reverse) ordered from Integrated DNA Technologies (San Diego, CA). The 232 bp PCR products were run on an agarose gel to confirm their size and purity. After gel extraction using the QIAquick Gel Extraction Kit (Qiagen), DNA concentration was measured with NanoDrop One C (Thermo Scientific) and normalized to 20 ng/μl. Samples were sent to GENEWIZ (New Jersey, USA) for next-generation sequencing (Amplicon-EZ) using an Illumina MiSeq platform and 250-bp paired-end reads. On average 400,000 reads were sequenced for each round. Sequencing data was processed using Galaxy^8^ with a cleanup protocol of: *fastp*^9^ for joining, trimming and quality filtering, and *transeq* for converting nucleotides to amino acids for further analysis.

We utilized FASTAptameR 2.0 to analyze the fully processed FASTA sequences from each round.^10^ FASTAptameR 2.0 enables one to count the number of reads for each sequence, determine the fold enrichment of each sequence over each round, cluster the sequences based on sequence similarity, and discover rate motifs and assess enrichment of those motifs specifically. To make the sequence logos of the sequencing populations, we used Logomaker.^11^ Sequence alignments with OAA homologs were done by BLAST^12^ and visualized using Jalview.^13^

To estimate the total diversity at each round from our undersampled amplicon sequencing, we used the Chao1 equation.^14^ Where $S_{est}$ is the estimated number of unique sequences, $S_{obs}$ is the number of observed unique sequences, $F_{1}^{2}$ is the number of single read sequences to the second power, and $2F_{2}$ is the number of double read sequences multiplied by 2:

$$S_{est}=S_{obs}+\frac{F_{1}^{2}}{2F_{2}}$$

**Variant Screen Cloning, Expression, and Purification**

Genes of variants V1-V10, and PM1-PM25 were ordered from GenScript. Inserts were cloned into pET-28b vectors (6xHis-TEV on the N-terminus) provided by GenScript using NEBuilder® HiFi DNA Assembly Master Mix and transformed into DH5alpha competent cells (Fisher). Plasmids were purified from DH5alpha cells using Monarch Plasmid Miniprep Kit (New England Biolabs) and transformed into BL21 (DE3) (New England Biolabs) competent cells. A single colony was picked from each plasmid transformation, then grown in LB broth (Fisher) overnight at 37 °C and finally stored at -80 °C in 25% glycerol (Promega). For protein expression, each variant was grown in 5 mL of LB broth + carbenicillin (G-Biosciences) at 37 °C/200 RPM. Upon reaching OD_600_ = ~0.6, 1 mM IPTG (Fisher) was added. After induction by IPTG, cultures were moved to 18 °C to shake at 200 RPM for ~16 hours. After expression was complete, cultures were spun down at 4,000 x g for 10 minutes. Cell pellets were resuspended in 100 µL NEBExpress® E. coli Lysis Reagent then frozen at -80 °C for 5 minutes. The sample was then thawed by rotating at room-temperature for 20 minutes. Samples were spun at 10,000 x g for 20 minutes to remove cell debris, the supernatant was transferred to new tubes that contained pre-washed Ni-charged MagBeads (GenScript). The cell lysate was rotated with beads for 30 minutes, then the placed on a magnetic rack for easy removal of supernatant. The beads were washed twice with 1x PBS (Fisher) + 20 mM imidazole (Fisher), then incubated in 100 μl 1x PBS + 250 mM imidazole for 10 minutes. Samples were placed on the magnetic rack and the supernatant was removed. Finally, the concentrations were estimated by A280 absorbance using Nanodrop One C. At this point, the samples were >95% in protein content by SDS-PAGE, with A260/A280 ratios under 0.8.

**Large-Scale Protein Expression and Purification**

For crystallography, NMR, BLI titrations, and pulldowns, proceeded with the following. Briefly, E. coli BL21(DE3) were transformed with a pET28b+ plasmid containing a N-terminal 6x-his tag followed by a TEV cleavage site and the OAA variant sequence. Cells were grown to an OD_600_ of 0.6 and induced with 1 mM IPTG at 18 °C for 16 h of shaking at 200 RPM. Cells were harvested by centrifugation at 4,000 x g, the supernatant was discarded, and the pellet was resuspended into wash buffer (1x PBS + 200mM NaCl, 20mM Imidazole pH 7.4). Cells were lysed via sonication (amplitude 60, 30s on with 45s rest for 6 min). The lysate was centrifuged at 25,000 x g for 25 minutes and the supernatant was filtered through a 0.2 µM filter. The clarified lysate was loaded on a HisTrap HP column and eluted with an imidazole gradient (5-100% imidazole over 10 CVs, where 100% is 500 mM). Small aliquots were removed at this stage for RNase B pulldowns and Ni-NTA Biosensor usage. The remaining Ni-NTA eluate was dialyzed into TEV cut buffer (1x PBS + 200mM NaCl + 10 mM beta-mercaptoethanol) and a 1:10 molar ratio of 6xHis-TEV protease:OAA was added to the sample during dialysis at 4 °C. The sample was applied over the HisTrap HP column again to capture all uncleaved protein and TEV protease, the flowthrough was collected in 1x PBS and concentrated for storage at -80 °C in 10 % glycerol. The purity and identity was established via SDS-PAGE. For crystallography, the sample was further purified by size-exclusion chromatography using a S75 16/60 column in 1x PBS with an ÄKTA Go protein purification system (GE Healthcare Life Sciences/Cytiva). These samples were then concentrated for storage at -80 °C in 10 % glycerol.

**SARS-CoV-2 Spike Expression and Purification**

SARS-CoV-2 Spike HexaPro variant was expressed using a mammalian expression system. Briefly, HEK293 cells were passaged to a final density of 0.6 - 0.7 x 10^6^ cells/mL before being transfected with a pαH plasmid (Addgene plasmid #299283 ; https://www.addgene.org/browse/sequence/299283) containing SARS-CoV-2 S HexaPro with a C-terminal 6x-his tag and a streptavidin tag. After 24 h, valproic acid (VPA) was added to the cells at a final concentration of 2.2 mM. Cells were harvested after 65-96 h, by pelleting the cells by centrifugation at 2,000 x g for 30 min at 4 ˚C. Supernatant was filtered through a 0.45 µM filter before being loaded onto Ni-NTA resin that has been equilibrated with wash buffer (20 mM sodium phosphate pH 7.4, 150 mM NaCl, and 30 mM imidazole). Ni-NTA resin is washed with 50 mL of wash buffer to remove nonspecific proteins. The supernatant is then incubated with the Ni-NTA resin and elution buffer (20 mM sodium phosphate pH 7.4, 150 mM NaCl, and 200 mM imidazole) for 30 min before being eluted. An SDS-PAGE gel is then run to evaluate which fractions to collect. Protein is then concentrated using a 100 kDa molecular weight cut off to ~1mg/mL before being loaded onto a Superose6 column in 1X PBS buffer using an ÄKTA Go protein purification system (GE Healthcare Life Sciences/Cytiva). Size exclusion fractions were then collected, evaluated by SDS-PAGE, concentrated and used immediately for BLI experiments or stored at -80 °C.

**Biolayer interferometry (BLI)**

BLI was conducted at 25 °C using an Octet® R2 (Sartorius) instrument coupled with Streptavadin biosensors for biotin-glycan and Ni-NTA biosensors for 6xHis-tagged proteins. Experimental buffer was 1x PBS + 0.1% BSA + 0.02% Tween 20. Ligand was loaded onto the biosensors followed by a baseline then association and dissociation phase. Biosensors were then regenerated in 10 mM Glycine pH 2.0. This process was repeated for varying concentrations of analyte. Data was processed in Octet analysis studio 13.0. For streptavidin biosensors, the samples were referenced to wells with no ligand. For Ni-NTA sensors they were double referenced by also referencing a well containing no analyte. The trace responses were aligned to the average of the final 5s of the baseline and inter-step corrected to the baseline. A Savitzky-Golay filter was applied to reduce high frequency noise. The association and dissociation phases were fit globally to a 1:1 single exponential binding model. Global fits had at minimum 4 concentrations for fitting. Extracted binding data was averaged over the replicates and reported with the average ± the standard deviation.

**RNase B Pulldowns**

OAA variants were purified with their 6xHis and 0.2 mg of protein was loaded onto Ni-charged Magbeads (GenScript) with a 10-minute incubation. The beads were then washed once with a 1x PBS + 20 mM imidazole buffer. 0.2 mg of RNase B was added to the beads in a 1x PBS buffer and incubated for 20 minutes. The beads were again washed once with a 1x PBS + 20 mM imidazole buffer and finally eluted with 1x PBS + 250 mM imidazole. The eluate was used directly for liquid chromatography coupled mass spectrometry (LC-MS). LC-ESI-TOF-MS was conducted on Agilent 6230 Accurate TOF-MS at the MMSF at UC San Diego.

**SARS-CoV-2 Neutralization Assays**

SARS-CoV-2 isolate “New York-PV08410/2020” (PANGO lineage B.1, BEI # NR-53514) was propagated on TMPRSS2-VeroE6 cells and titered by fluorescent focus assay on TMPRSS2-VeroE6 cells. Neutralization assays were performed as previously described with modifications.^15^ TMPRSS2-VeroE6 cells were seeded in 96-wells plates at 15,000 cells per well the day before infection. OAA variants were threefold serially diluted in OptiMEM and incubated with approximately 100 focus-forming units of SARS-CoV-2 in technical triplicate for 1 hour at 37 °C, then 35µl of virus OAA mixtures was transferred to cells. After 1 hour infection at 37 °C, 100µl per well of viscous overlay (1% methylcellulose in MEM supplemented with 2% FBS (Biowest), 1x penicillin/streptomycin, 1x non-essential amino acids, 1x GlutaMAX, and 10mM HEPES) was added to restrict viral spread to neighboring cells. Plates were fixed with 4% formaldehyde after 24 hours of growth, and foci were visualized by immunofluorescence using anti-nucleocapsid antibody (GeneTex, gtx135357) with AlexaFluor 555 anti-rabbit secondary antibody and Hoechst 33342 nuclear counterstain. Whole well images were captured with a 4x objective in a Cytation 5 plate imager (Agilent), and spots were automatically counted in stitched images with the associated Gen5 software. Foci counts were normalized to 3-6 media-only infected wells on each plate and expressed as percent infectivity. Figures were arranged in plate layout using ImageJ Stitching plugin.^16^

**Protein Crystallography**

To determine crystallization conditions for our variant V4V4, we sent purified sample to the National Crystallization Center (Buffalo, NY, USA) for high throughput screening against 1536 conditions.^17^ From the 1536 condition screen, we selected 0.1 M KNO_3_ (VWR), 0.1 M BIS-TRIS propane (MP Biomedicals) pH 7.0, 80% PEG 400 (Thermo Scientific) as the optimal condition. We proceeded with crystallization using the hanging-drop method. V4V4 was first buffer exchanged into 20 mM Tris pH 8.0, 100 mM NaCl, 0.03% NaN3, and 3α,6α-mannopentaose was dissolved in MQ H_2_O. To co-crystallize we added 1 μl of V4V4 (20 mg/mL), 1 μl of 3α,6α-mannopentaose (5.4 mM) to give a 1:4 ratio of protein:ligand, and 2 μl of 0.5x diluted crystallization buffer. 1 mL crystallization buffer at a 0.5x dilution was added below the hanging drop, the well was sealed off with vacuum grease and left to crystallize at 25 °C, where crystals formed after several weeks. Crystals were looped with 10% glycerol as cryoprotectant and flash frozen in liquid nitrogen for delivery to the Advanced Light Source.

Diffraction data were collected at beamline 24ID-C at the Advanced Light Source at Argonne National Laboratory (beamtime award DOI: https://doi.org/10.46936/APS-191148/60014907). Data were processed by the RAPD2 pipeline (https://git.nec.aps.anl.gov/rapd/rapd), which uses XDS^18^ for indexing and integration, AIMLESS^19^ for merging, and CTruncate^20^ for conversion to structure factors. The structure was determined by molecular replacement in PHASER^21^ using (PDB 3S5X), manually rebuilt in COOT,^22^ and refined in phenix.refine^23^ using individual atomic position and isotropic B-factor refinement, plus TLS refinement (one TLS group per chain). Structures were prepared for figures and analyzed using UCSF ChimeraX.^24^ The structure was deposited into the PDB with PDB code 10KB.

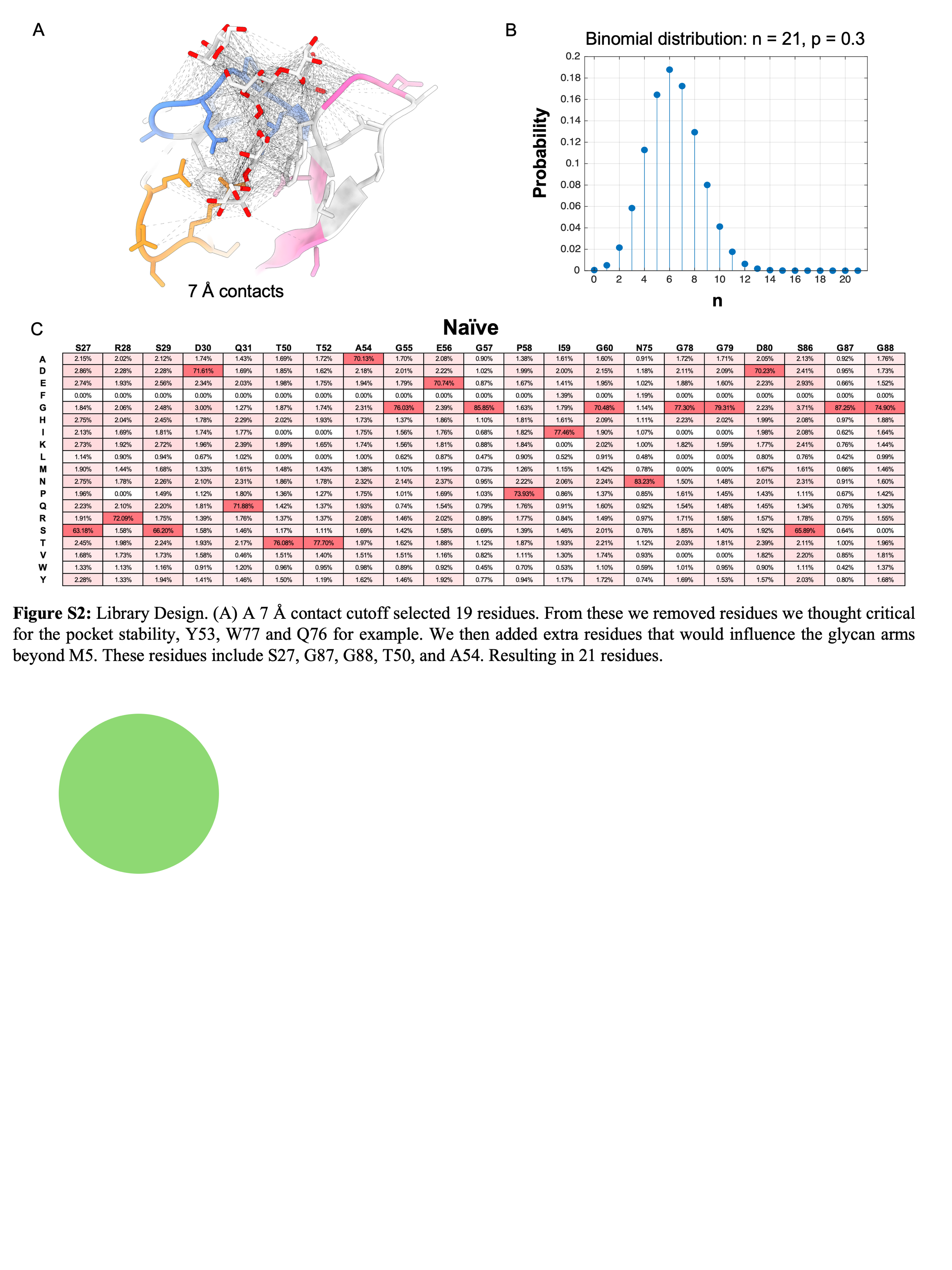
 **Supplemental Figure 1. Library design.**

(A) Using PDB:3S5X, a 7 Å contact map was made from the 3α,6α-mannopentaose ligand to select 19 WT OAA residues. From these, we removed residues we thought critical for the pocket stability, Y53, W77 and Q76 for example. We then added extra residues that could influence the glycan arms beyond M5. These residues include S27, G87, G88, T50, and A54 for a total of 21 residues. (B) To determine the mutation rate of our 21 residues, we aimed for a distribution that would still explore all of the single point mutation. We therefore aimed for a 30% chance of mutation at each site which results in a mean of 6.3 mutations per variant. (C) Heat map table showing the percentage of each amino acid at each residue site in the naïve library. Notable is our omission of cysteine and the infrequent use of phenylalanine because these are largely solvent exposed sites. Red signifies higher percentage, while white signifies 0%.

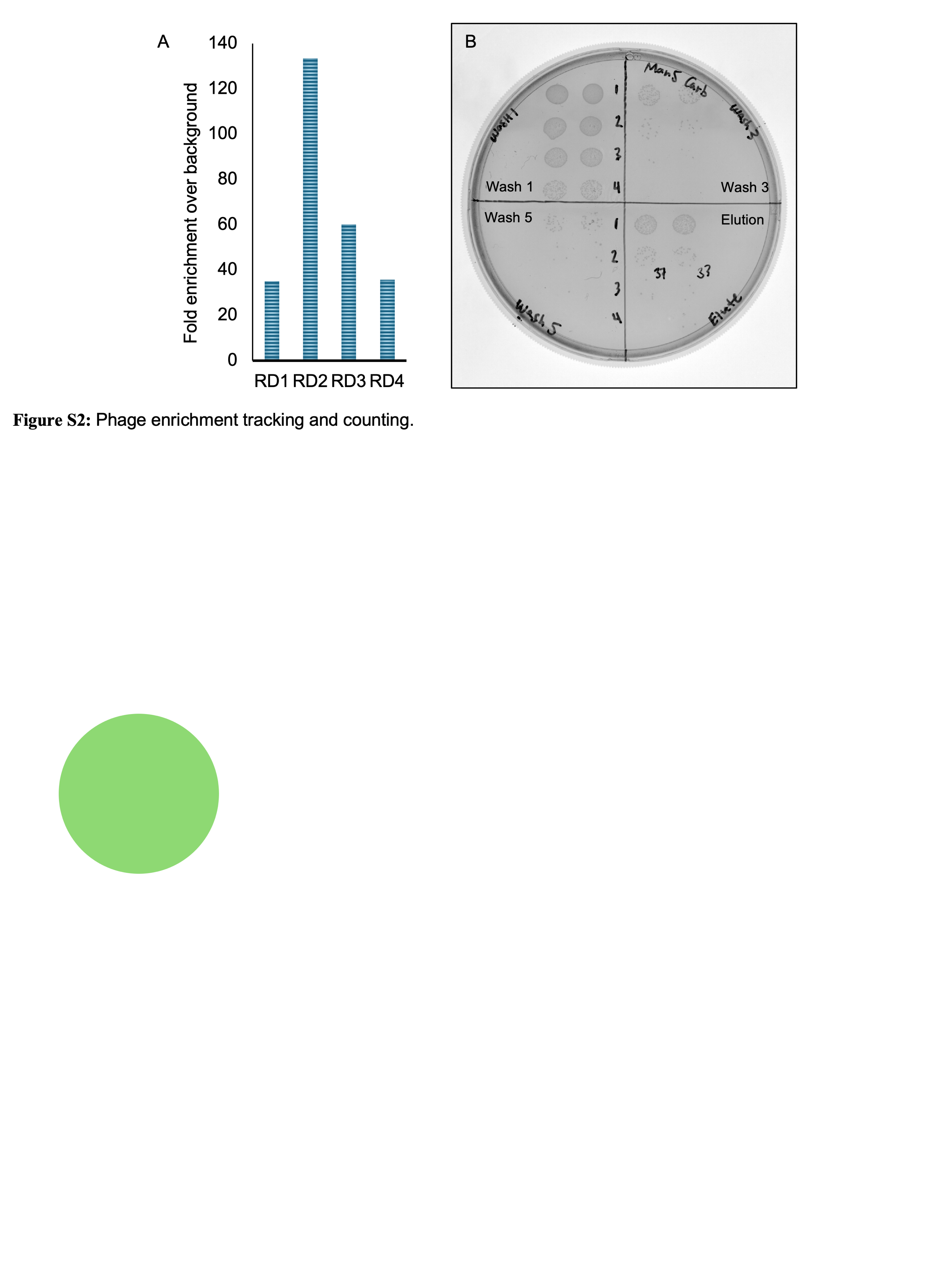
**Supplemental Figure 2. Phage enrichment and plate spots.**

(A) A bar graph showing the phage yield for biotin-M5-coated beads enriched over unbound strep-beads. Round 1 through Round 4 is shown. (B) An example of a phage titer on agar to count the yield of phages from a bead selection procedure. Shown are the titrations from 10^-1^ to 10^-4^ for wash 1, 3, 5, and the elution. Importantly, the phage count is >10x in the elution over the final wash step.

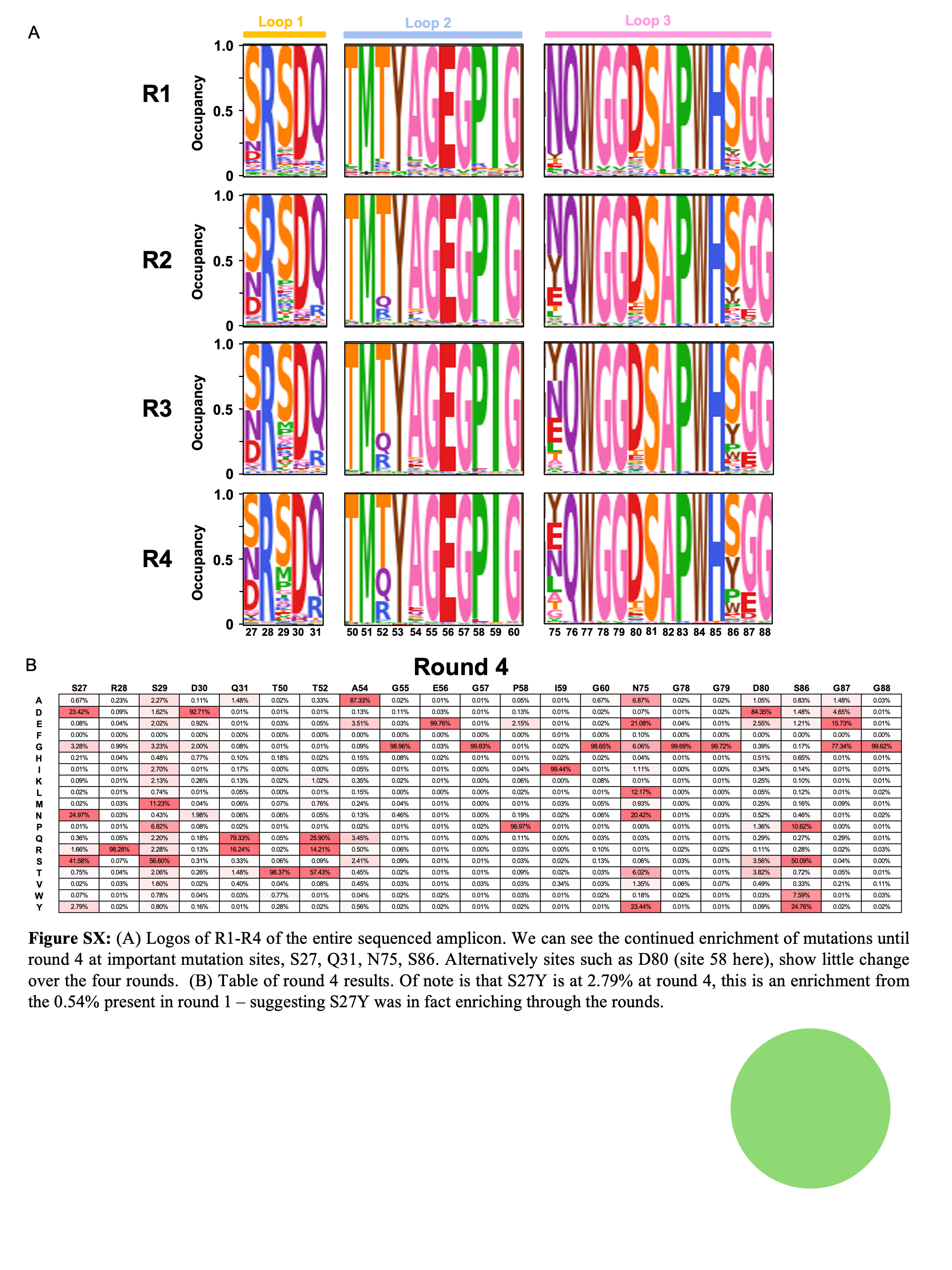
**Supplemental Figure 3. Round by round mutational presence.**

A) Sequence logos of round 1 to round 4 of the entire sequenced amplicon. We note a continued enrichment of mutations until round 4 at important mutation sites, S27, Q31, N75, S86. Alternatively, sites such as D80 (site 58 here), show little change over the four rounds. (B) Heat map table of round 4 results. S27Y is at 2.79% at round 4, enriching from 0.54% present in round 1 – suggesting S27Y was benefitting from continued rounds of selection. Red signifies higher percentage, while white signifies 0%.

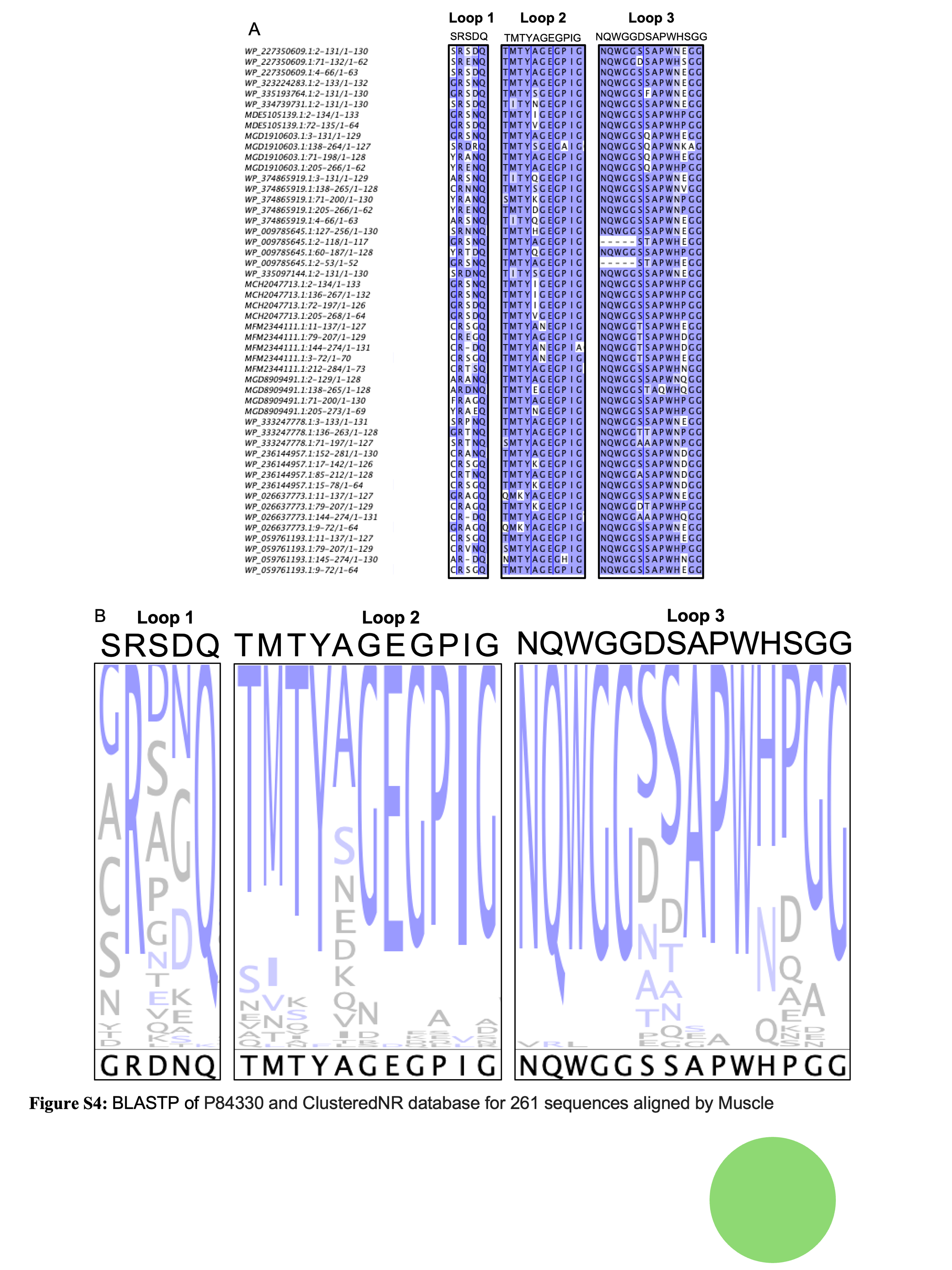

**Supplemental Figure 4. Sequence alignment of OAA family lectins.**

(A) BLASTP of P84330 and ClusteredNR database for 261 sequences aligned by Muscle. Top 40 sequences are shown for reference. (B) The WT OAA sequence used in the study is located on top of the sequence logo from sequence alignment. The consensus sequence is denoted on the bottom. Notable is the evolutionary lack of mutation at Q31 and N75, and no S86Y mutations – all important mutations in our study. We also see similarities such as the high mutational frequency in S27, S29, D30, and S86.

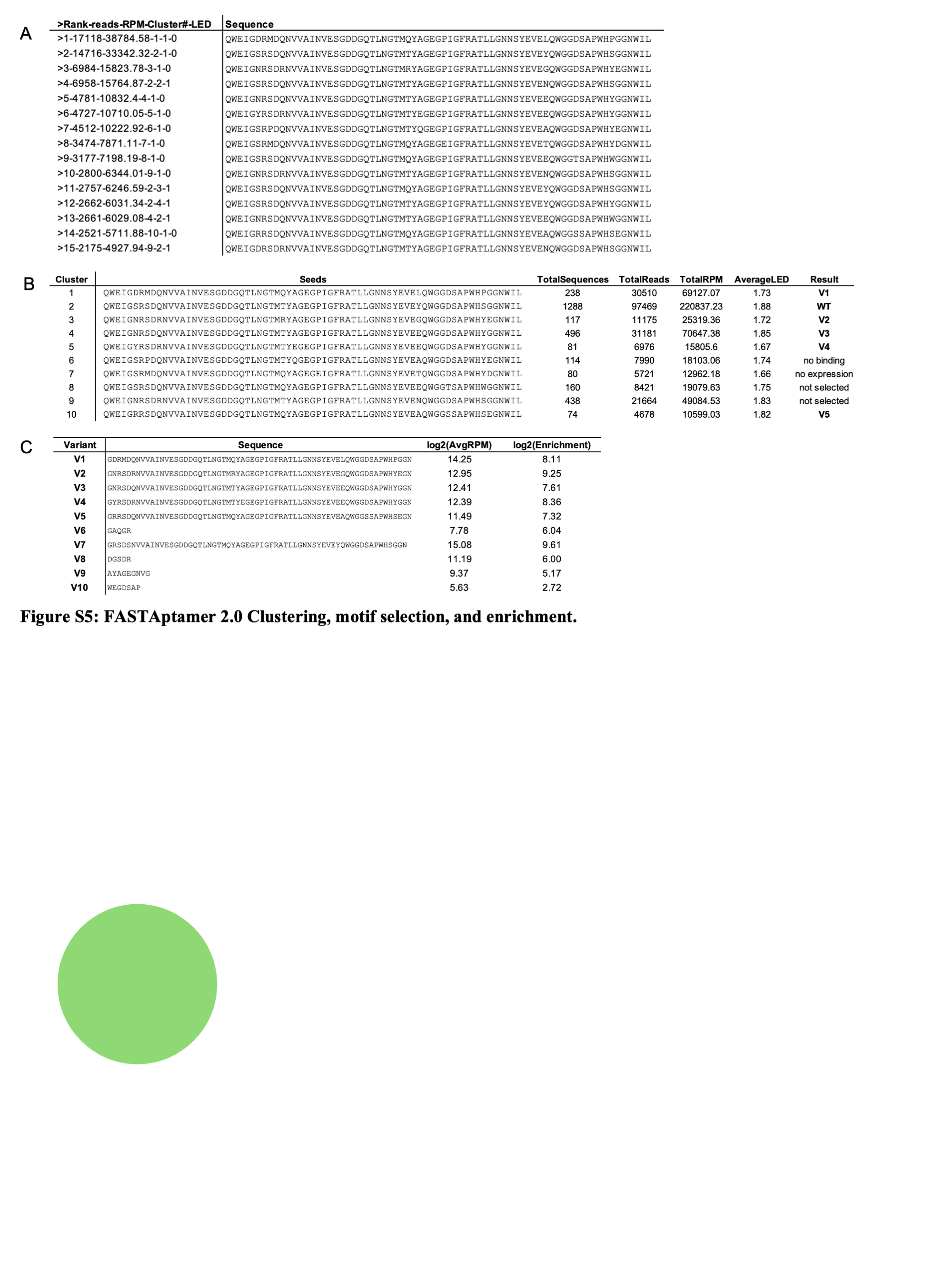
**Supplemental Figure 5. Round 4 enrichment clusters.**

(A) Example of top 15 read sequences entering clusters. Where RPM is reads per million, Cluster# is which cluster group they enter, and LED is the Levenshtein edit distance which describes how different the sequence is from the cluster seed. The clusters were set to have a max LED of 2, so if a sequence is more than 2 mutations different from the seed it gets screened against the next cluster. (B) The cluster data is shown, highlighting the number of unique sequences, total reads, and the resulting variant (V1-V5) made from the cluster seed sequence. (C) The volcano plot data for each variant (V1-V10) shown as log_2_(Average RPM)_R1+R4_ and log_2_(Enrichment)_R1:R4_. The motif derived variants (V6-V10) were analyzed for their unique motifs, rather than whole sequences.

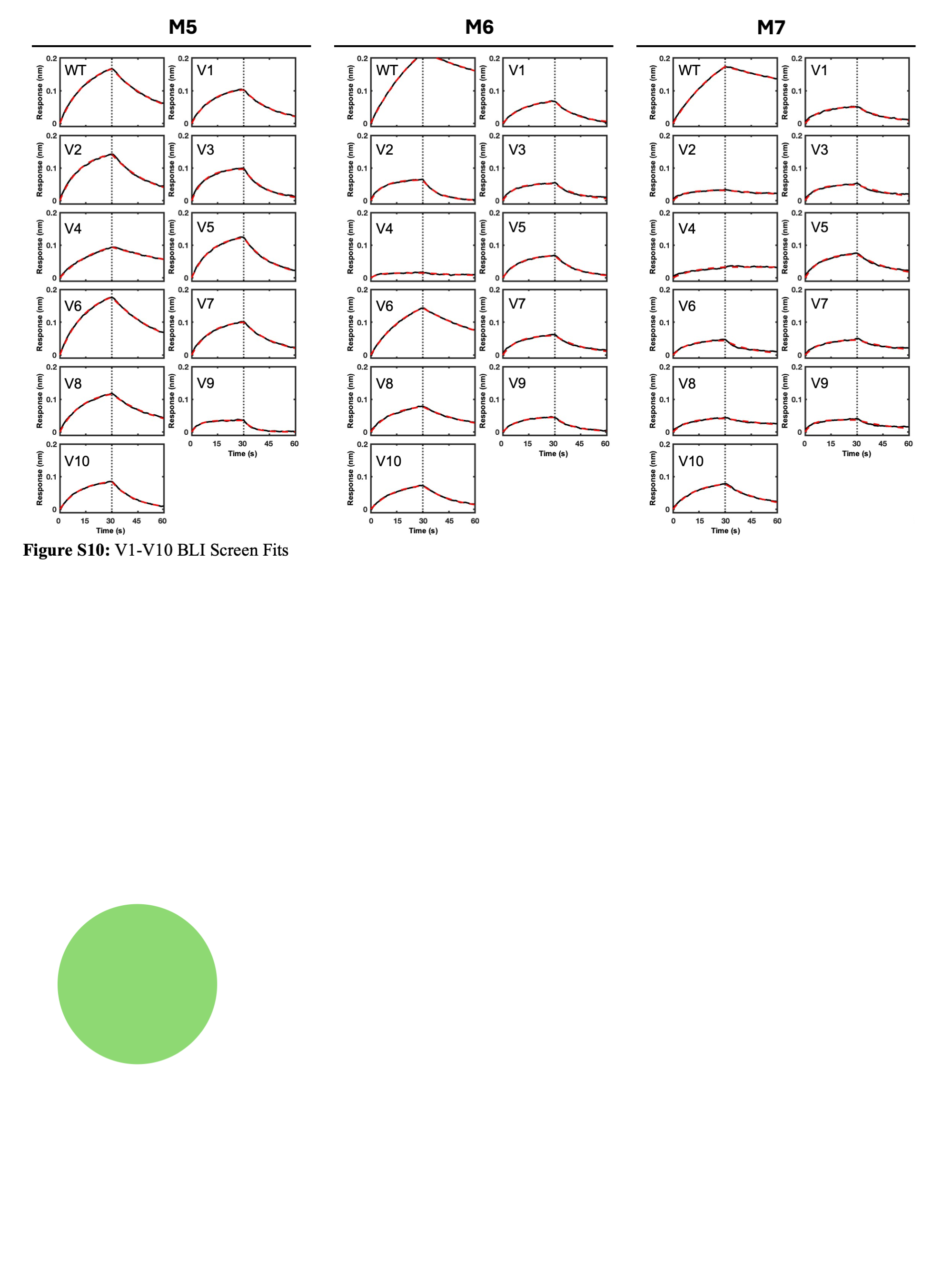
**Supplemental Figure 6. BLI fits of variant glycan screen.**

Fits are shown for V1-V10 and WT for M5 (*left)*, M6 (*middle)*, and M7 (*right*) from a single replicate. The complete fitting results can be found in Supplemental Data 2.

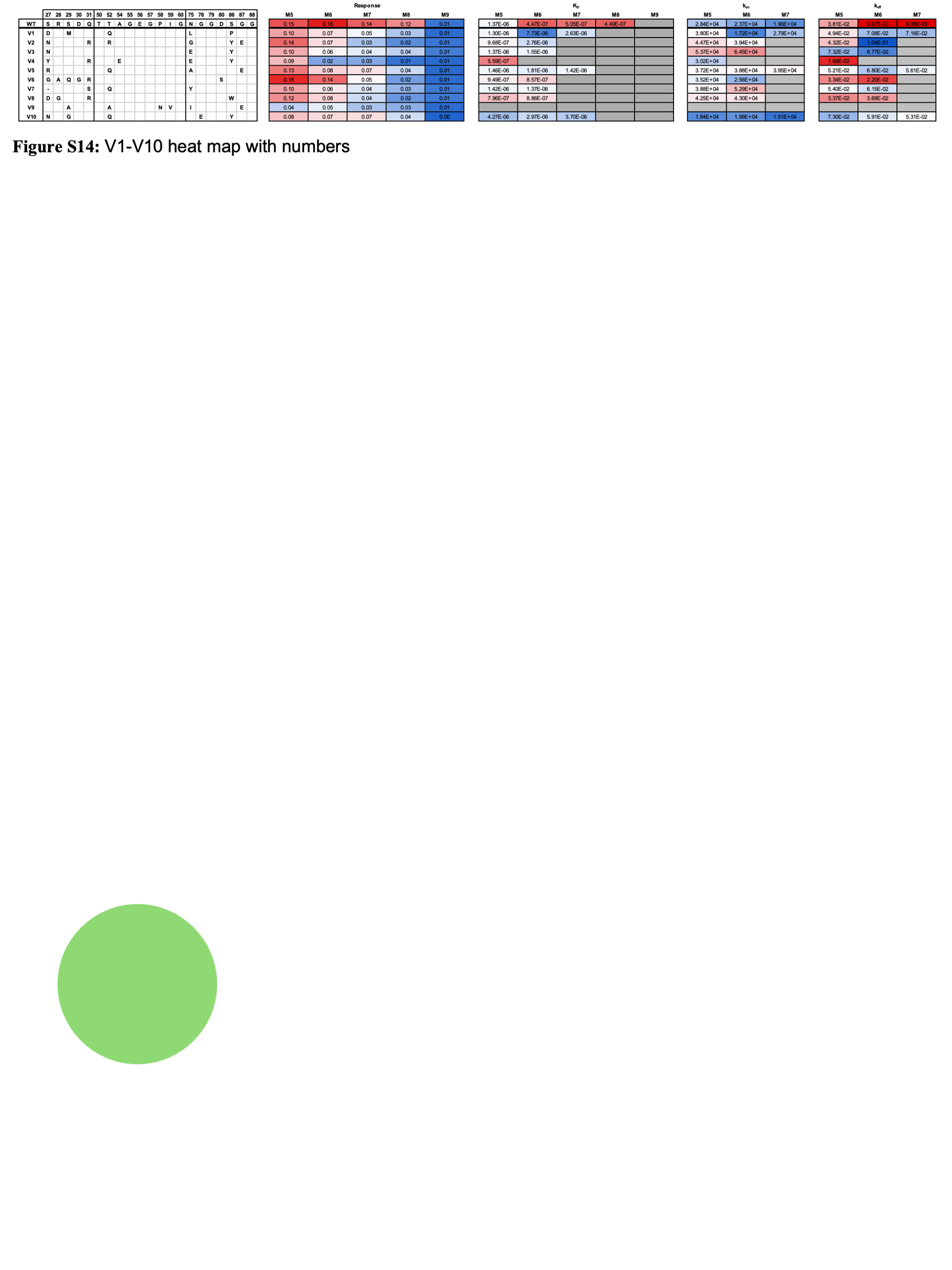
**Supplemental Figure 7. Heat map of variant screen with numbers.**

Heat map of variant mutations with residue numbers with data corresponding to Figure 3 in the main text. Each variant was screened at 1 μM against each biotin-glycan, M5-M9. Heat maps of BLI data are shown for sensor response, binding affinity (K_D_), association rate (k_on_), and dissociation rate (k_off_). Heat map color red signifies better binding character, while blue signifies worse binding character. Gray boxes mark variant-glycan interactions that did not reach 0.05 nm response and therefore were not fit for kinetics. The response heat map shows a response of 0 to 0.2 (*red*) nm with white at 0.05 nm to obviate the threshold of confidence. The K_D_ heat map shows a range of 0.1 (*red*) to 10 μM. The K_on_ heat map shows 1x10^4^ to 1x10^5^ (*red*) 1/Ms. The K_off_ heat map shows 1x10^-2^ (*red*) to 1x10^-1^ 1/s. The values were determined by the average value of a duplicated BLI screen.

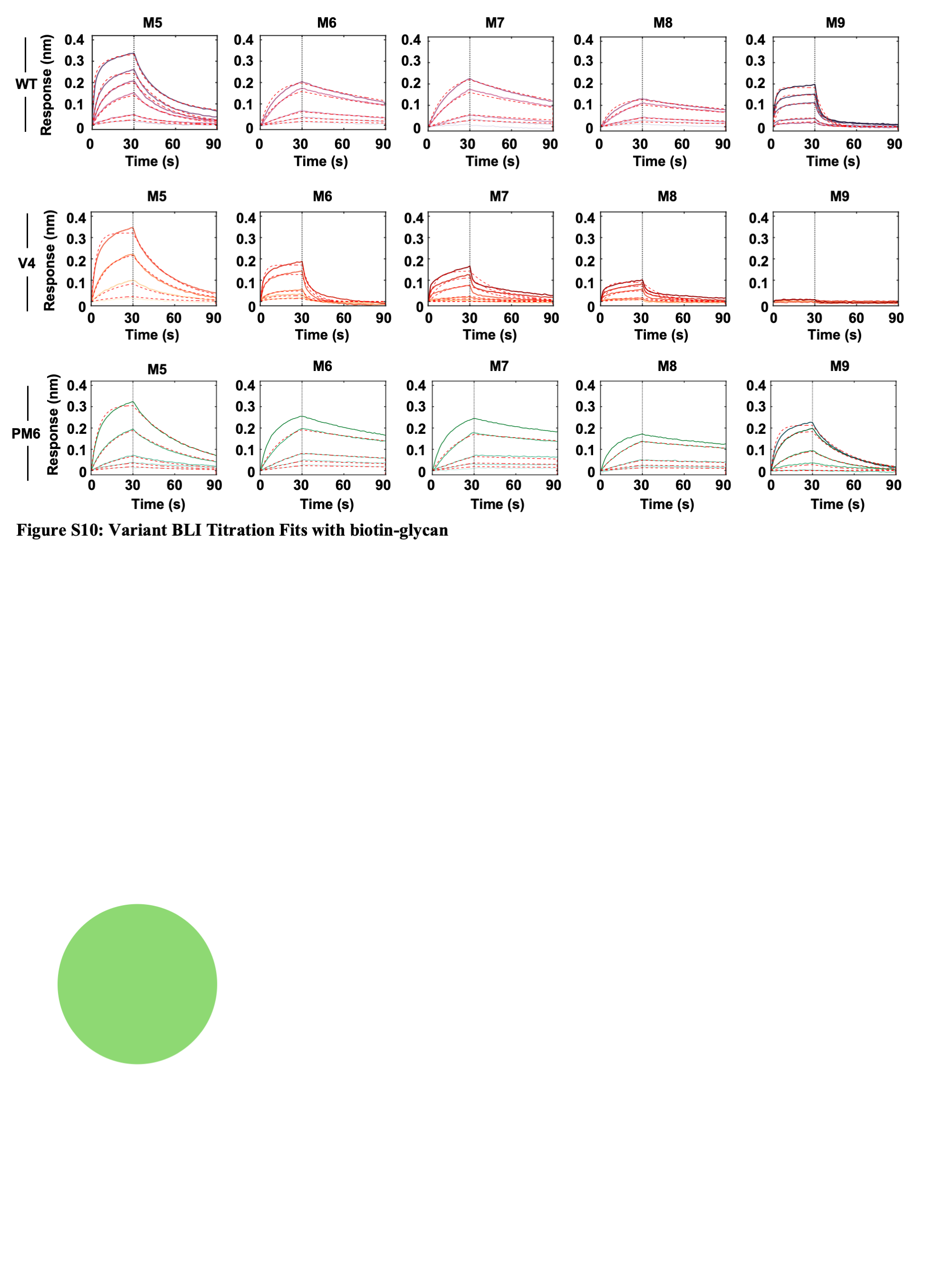
**Supplemental Figure 8. BLI fits for binding titrations of WT, V4, and PM6 across M5-M9 glycans.**

BLI fits of WT (*top*), V4 (*middle*), and PM6 (*bottom*) titrations, the data correspond to Figure 3 and Figure 5 in the main text. Complete fitting results can be found in the Supplemental Data 2.

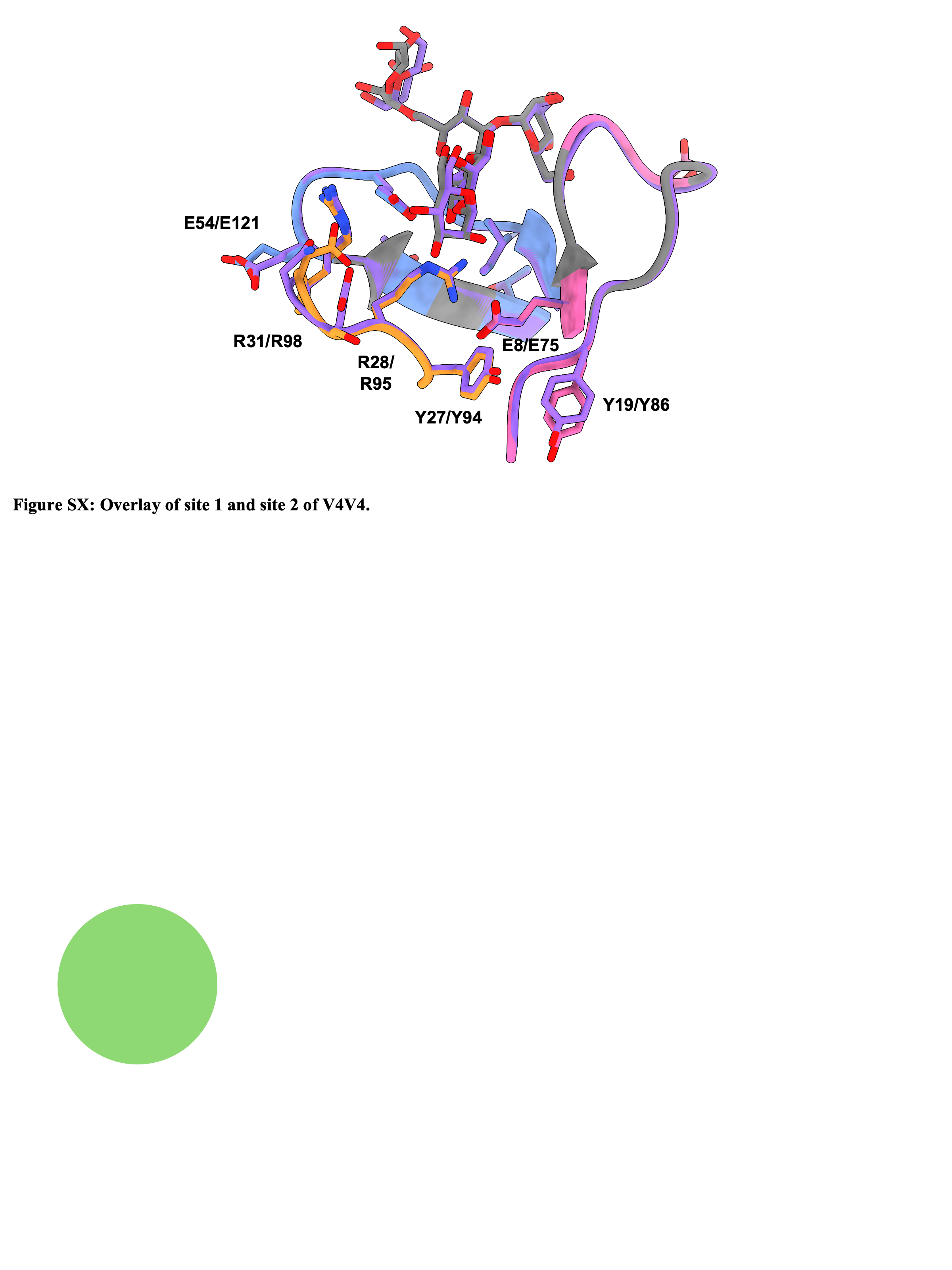

**Supplemental Figure 9. Overlay of site 1 and site 2 of V4V4.**

Overlay of V4V4 site 1 (gray, loop colors), and site 2 (purple), showing the overall Cα RMSD of 0.25 Å.

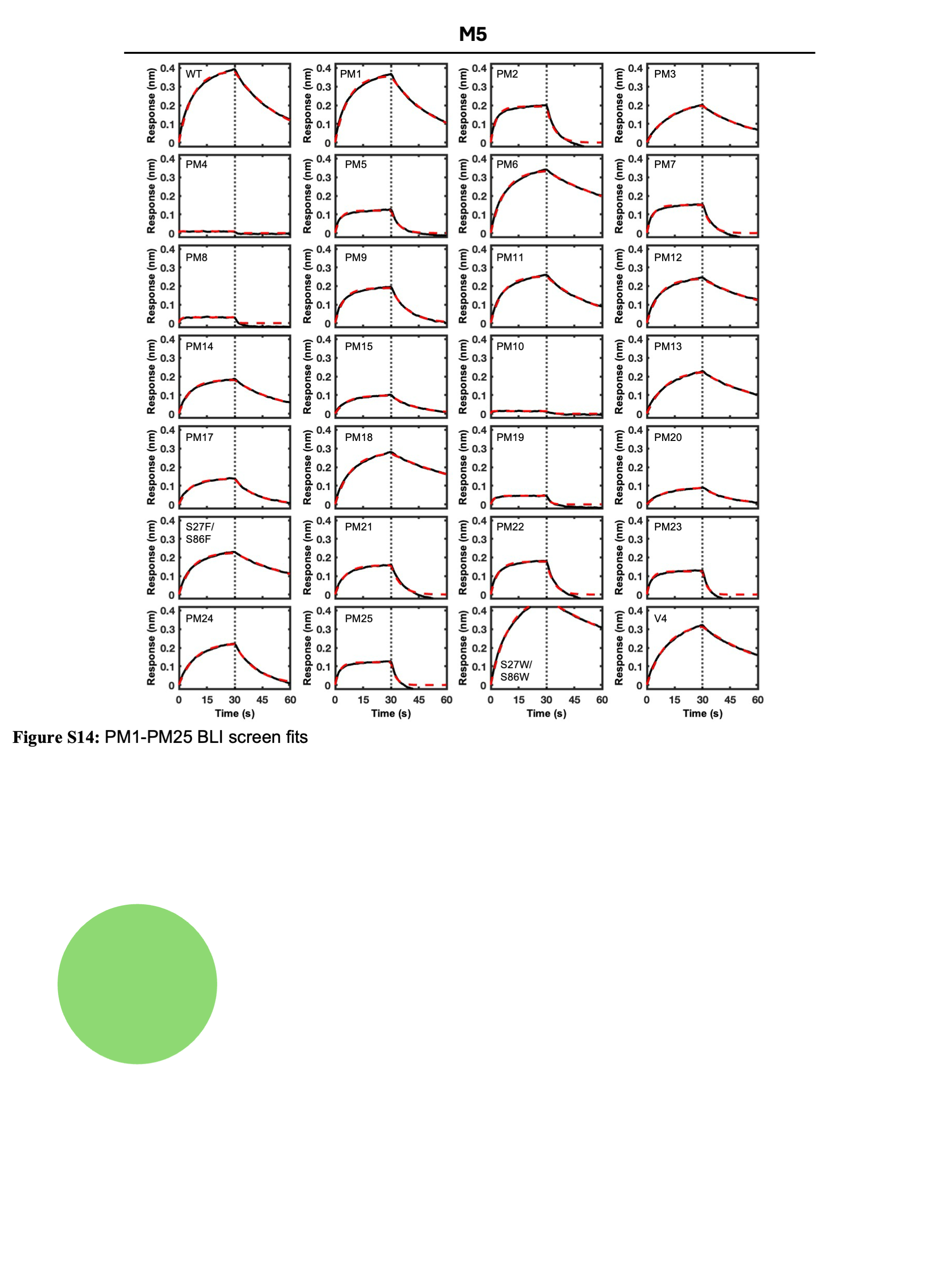
**Supplemental Figure 10. BLI fits of point mutation screen.**

Fits are shown for PM1-PM25, WT, V4, S27F/S86F, and S27W/S86W variants for M5 as an example from a single replicate. The complete fitting data can be found in the Supplemental Data 2.

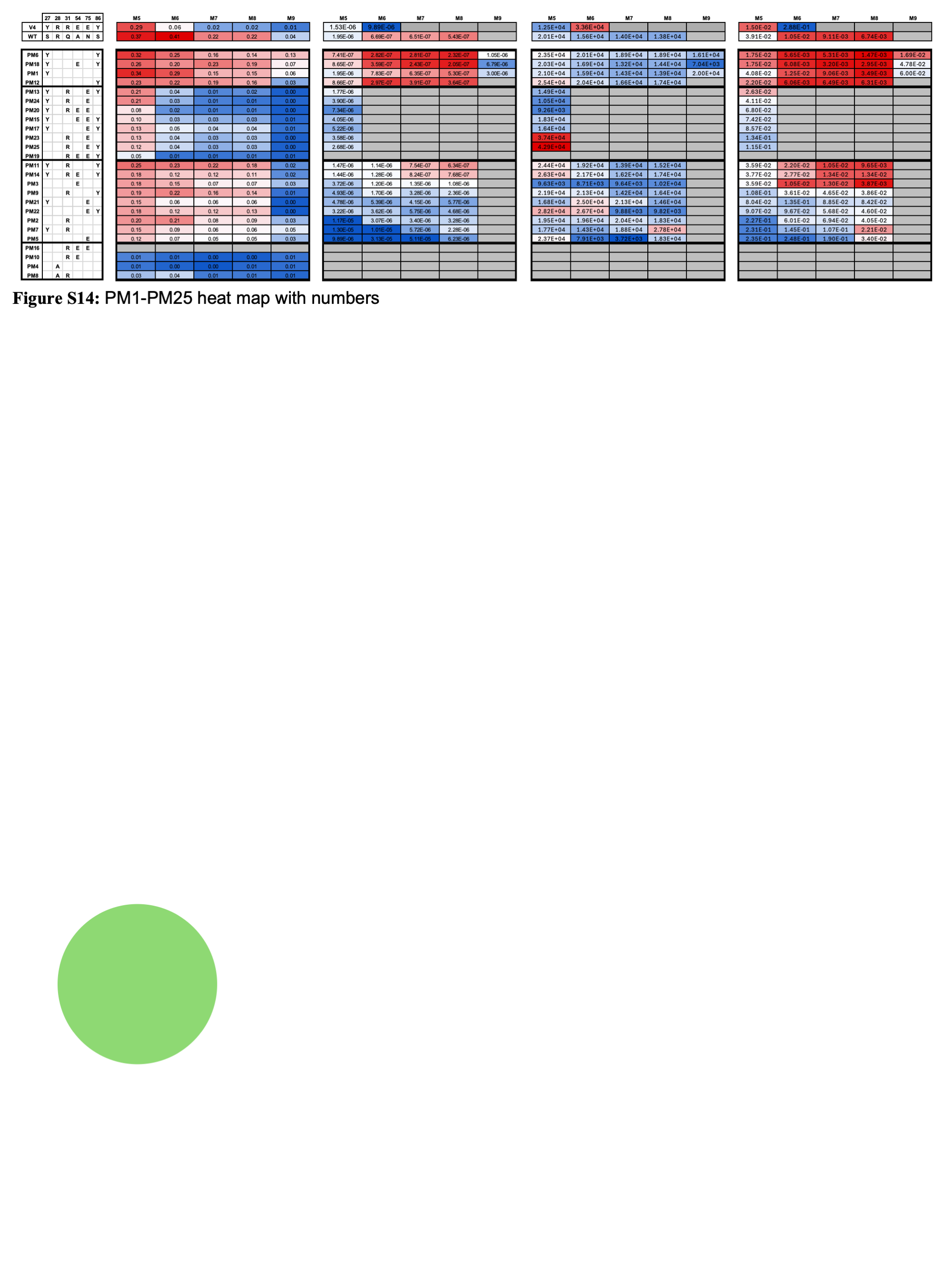
**Supplemental Figure 11. Heat map of point mutation screen with numbers.**

Heat map of V4-based point mutation variants with residue numbers corresponding to Figure 5 in the main text. Each variant was screened at 5 μM against each biotin-glycan, M5-M9. Heat maps of BLI data are shown for sensor response, binding affinity (K_D_), association rate (k_on_), and dissociation rate (k_off_). Heat map color red signifies better binding character, while blue signifies worse binding character. Gray boxes mark variant-glycan interactions that did not reach 0.05 nm response and therefore were not fit for kinetics. The response heat map shows a response of 0 to 0.4 (*red*) nm with white at 0.05 nm to obviate the threshold of confidence. The K_D_ heat map shows a range of 0.1 (*red*) to 10 μM. The K_on_ heat map shows 3x10^3^ to 4x10^4^ (*red*) 1/Ms. The K_off_ heat map shows 1x10^-3^ (*red*) to 3x10^-1^ 1/s. The values were determined by the average value of a duplicated BLI screen.

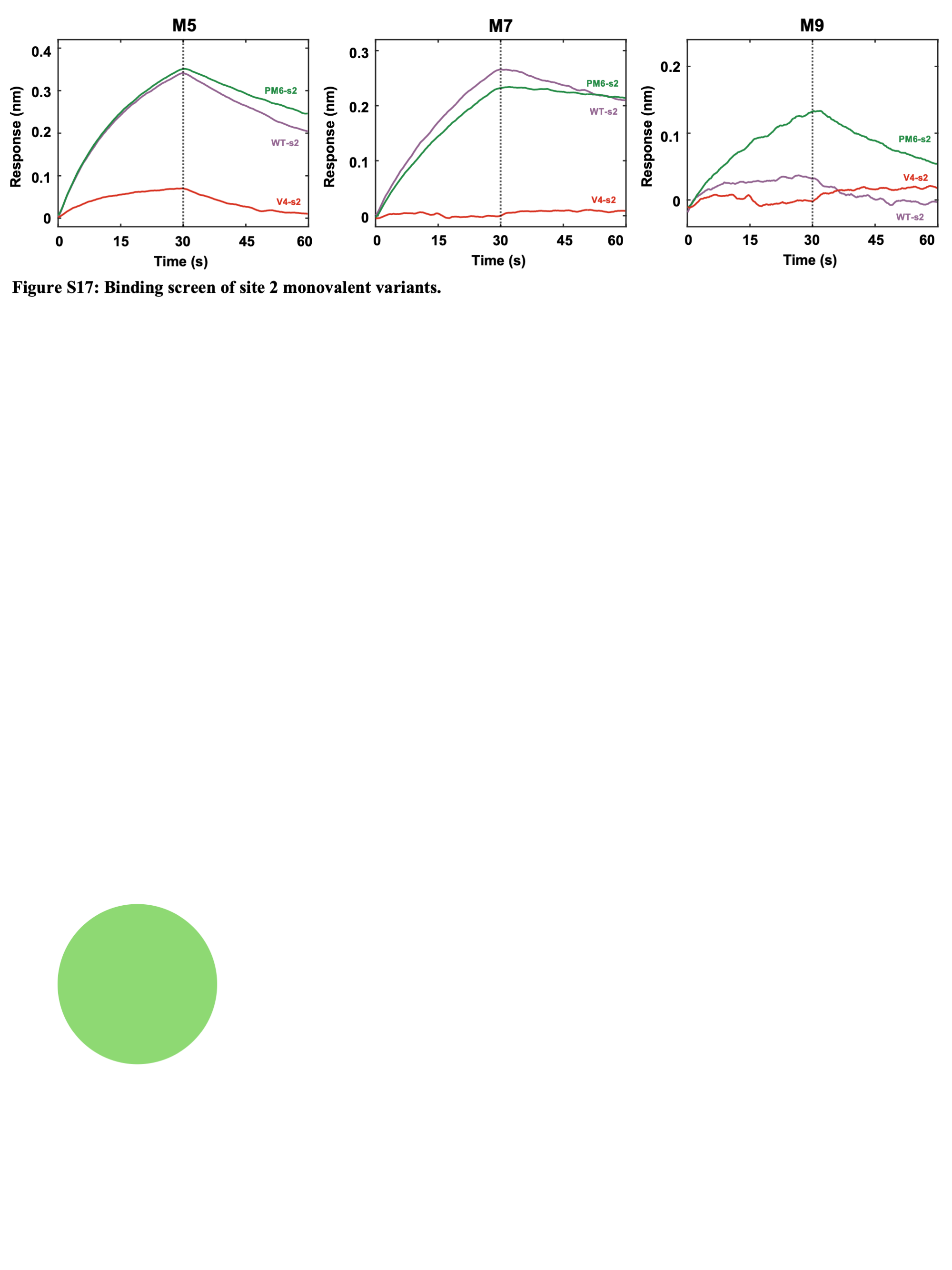
**Supplemental Figure 12. Binding screen of site-2 monovalent variants.**

WT-s2, V4-s2, and PM6-s2 are the discovered mutations grafted onto site 2 of OAA. The variants were screened against M5 (left), M7 (middle), and M9 (right) at 1 μM. This screen serves as a confirmation that the mutations follow the same trend when transferred to site 2.

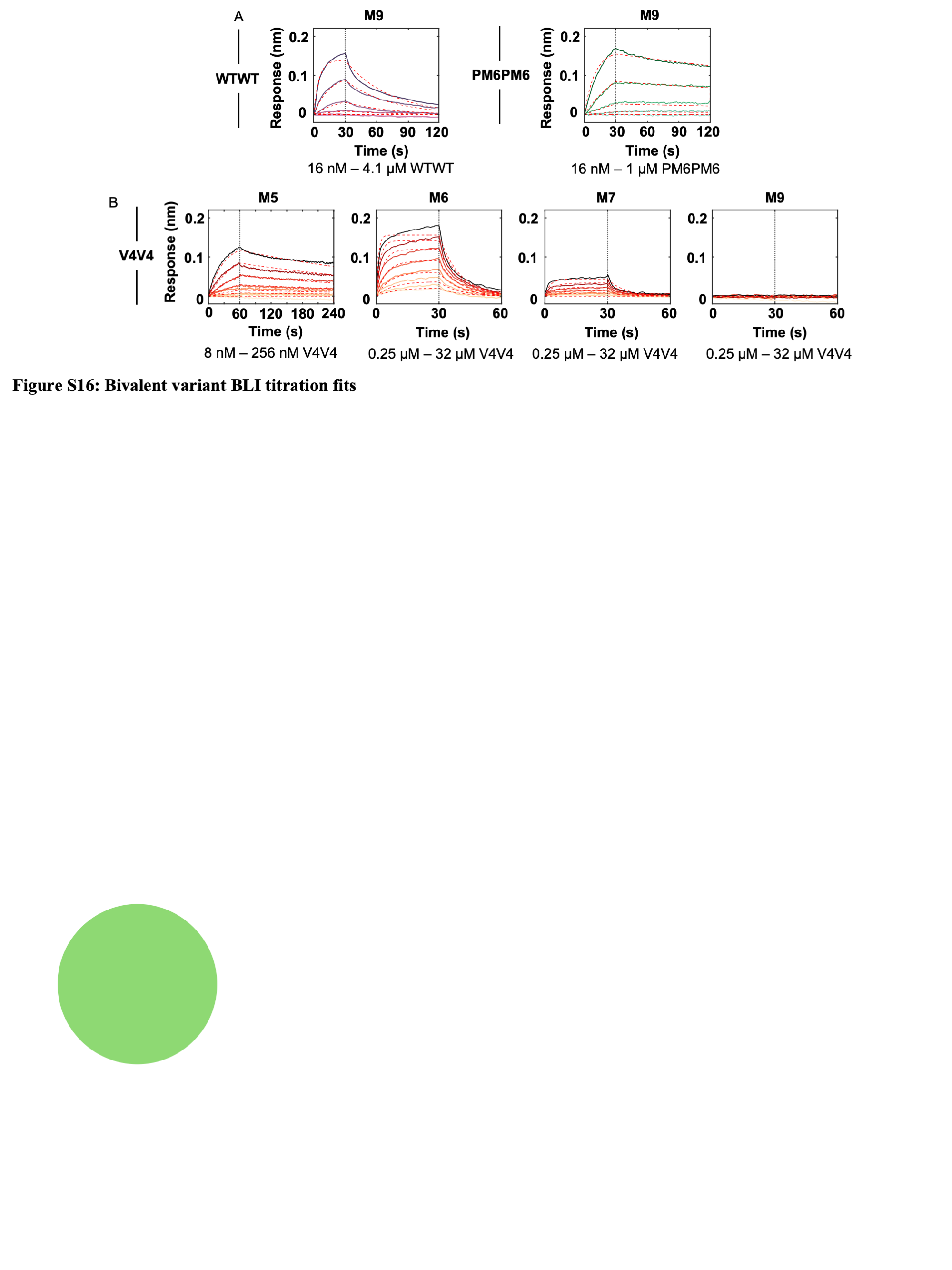
**Supplemental Figure 13. Bivalent variant BLI titration fits with biotin-glycans.**

(A) BLI fits for WTWT (left) and PM6PM6 (right) on M9. (B) BLI fits for V4V4 across M5, M6, M7, and M9. Complete fitting results are available in the Supplemental Data 2.

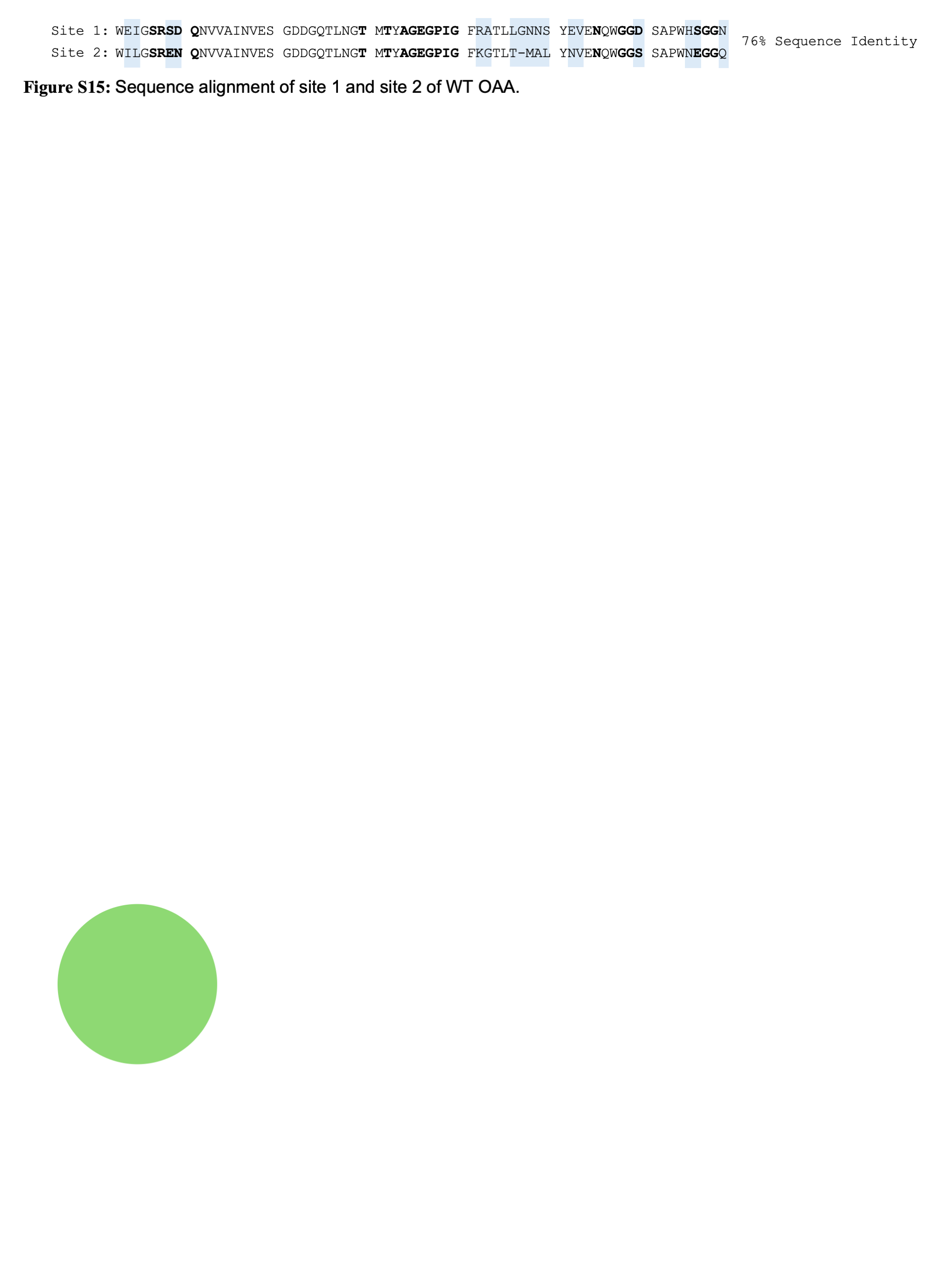

**Supplemental Figure 14. Sequence alignment of WT OAA site 1 and site 2.**

Sequence alignment of each side of the OAA beta-barrel showing only 76% sequence identity match. Bolded letters correspond to the residues used in the mutation library. Blue highlights signify mismatches.

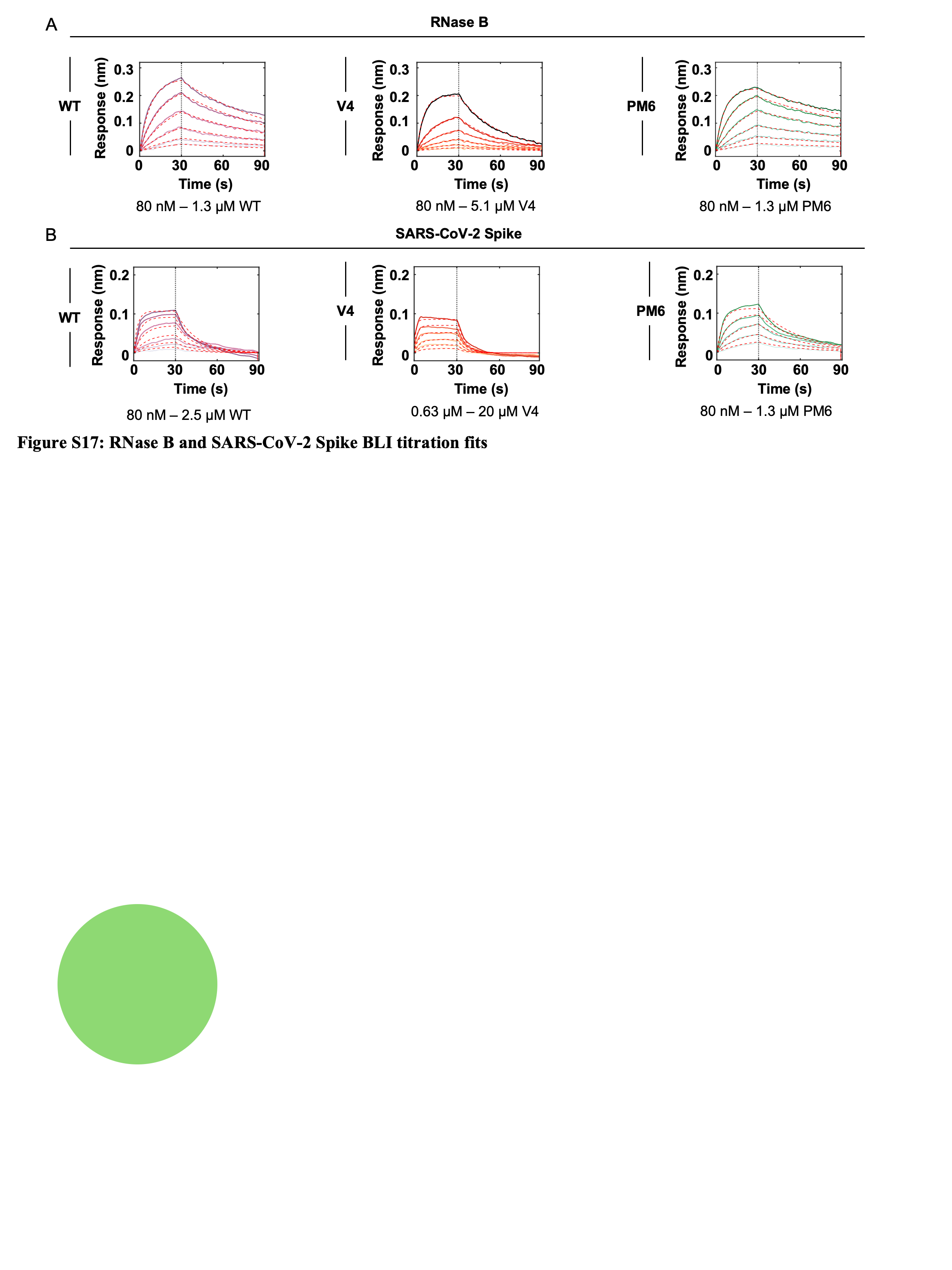

**Supplemental Figure 15. Glycoprotein BLI titration fits with monovalent variants.**

(A) Fits of 6xhis-tagged WT, V4, and PM6 against RNase B. (B) Fits of WT, V4, and PM6 against 6xhis-tagged SARS-CoV-2 Spike protein. Complete titration fitting results can be found in the Supplemental Data 2.

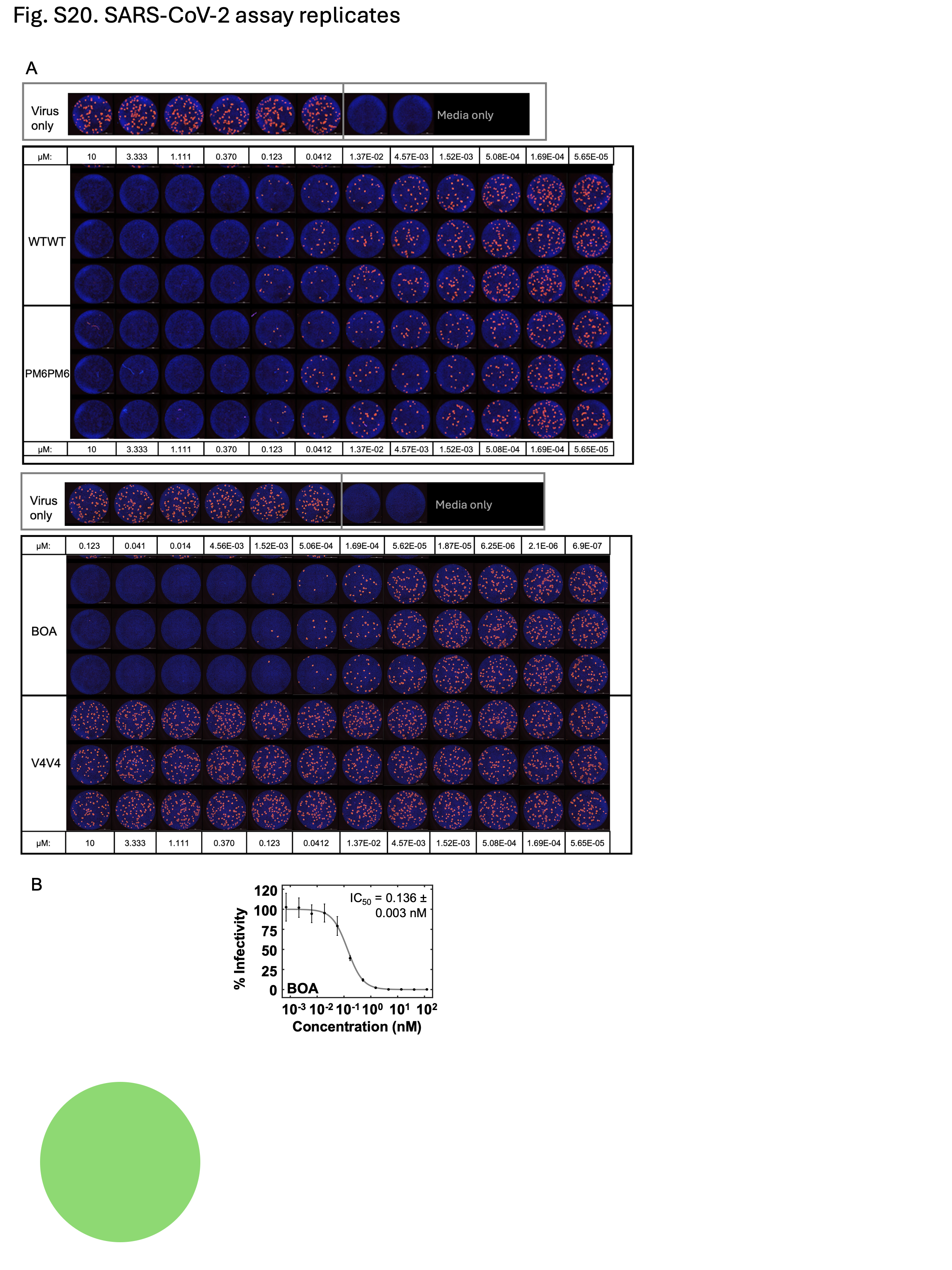

**Supplemental Figure 16. SARS-CoV-2 inhibition detection by focus reduction neutralization assay.**

(A) Representative example of neutralization assay. Foci of infection were counted by software in stitched images of whole wells of 96-well plates. Controls shown on top for normalization of all foci counts. Assay was repeated in n=3 independent experiments with triplicate wells per dilution. Images were arranged in plate layout using ImageJ Stitching plugin. (B) Concentration-dependent inhibition of SARS-CoV-2 lineage B.1 by BOA, as measured by neutralization assay, yielding an IC50 value of 0.136 ± 0.003 nM.

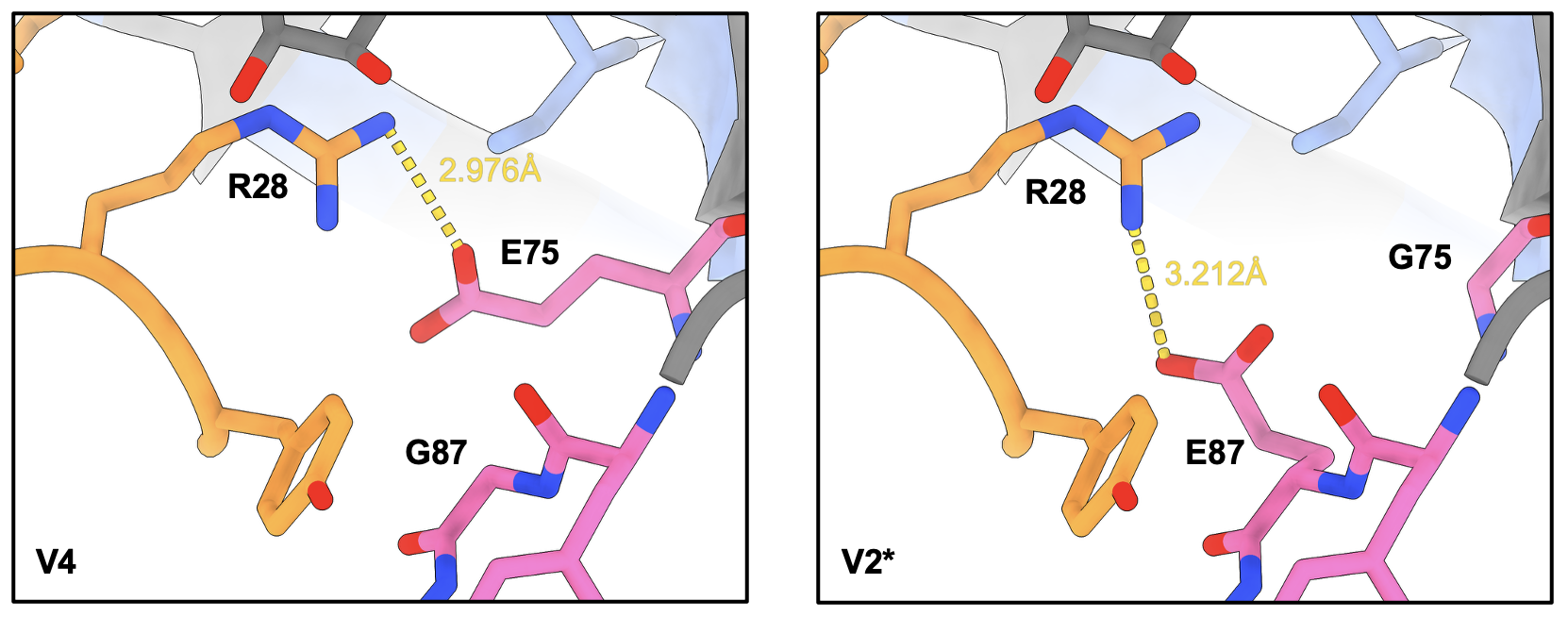

**Supplemental Figure 17. Possible mechanism of V2 selectivity for M5.**

Mutational change of V4 site from V4V4 crystal structure into V2*, a hypothetical position for the G87E mutation. As shown, the glutamate from site 87 can mimic the site 75 glutamate for the R28 rotamer flip.

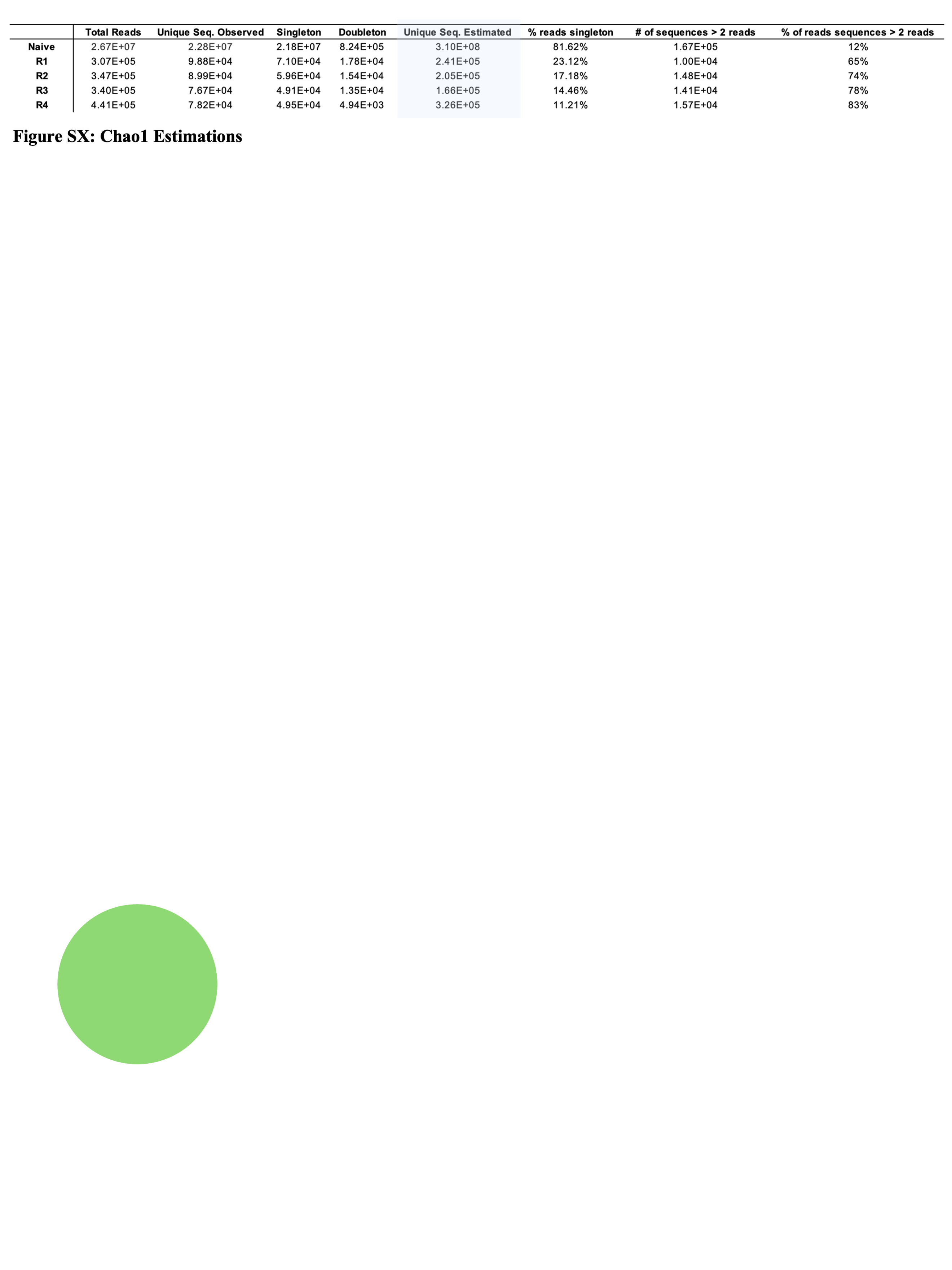

**Supplemental Table 1. Amplicon sequencing results and estimates.**

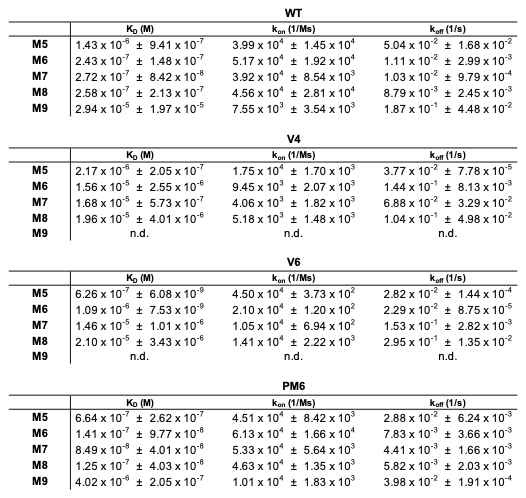

**Supplemental Table 2: Variant binding affinity and kinetics.**

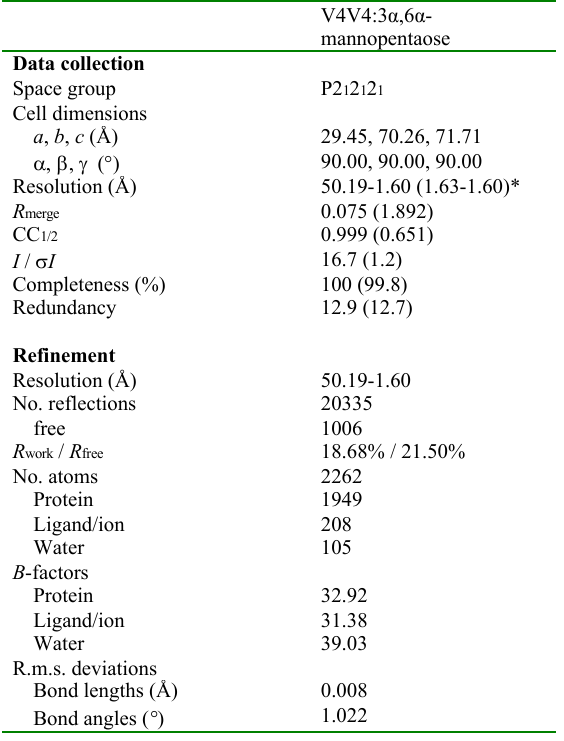

*Values in parentheses are for highest-resolution shell.

**Supplemental Table 3: Data collection and refinement statistics (molecular replacement)**.

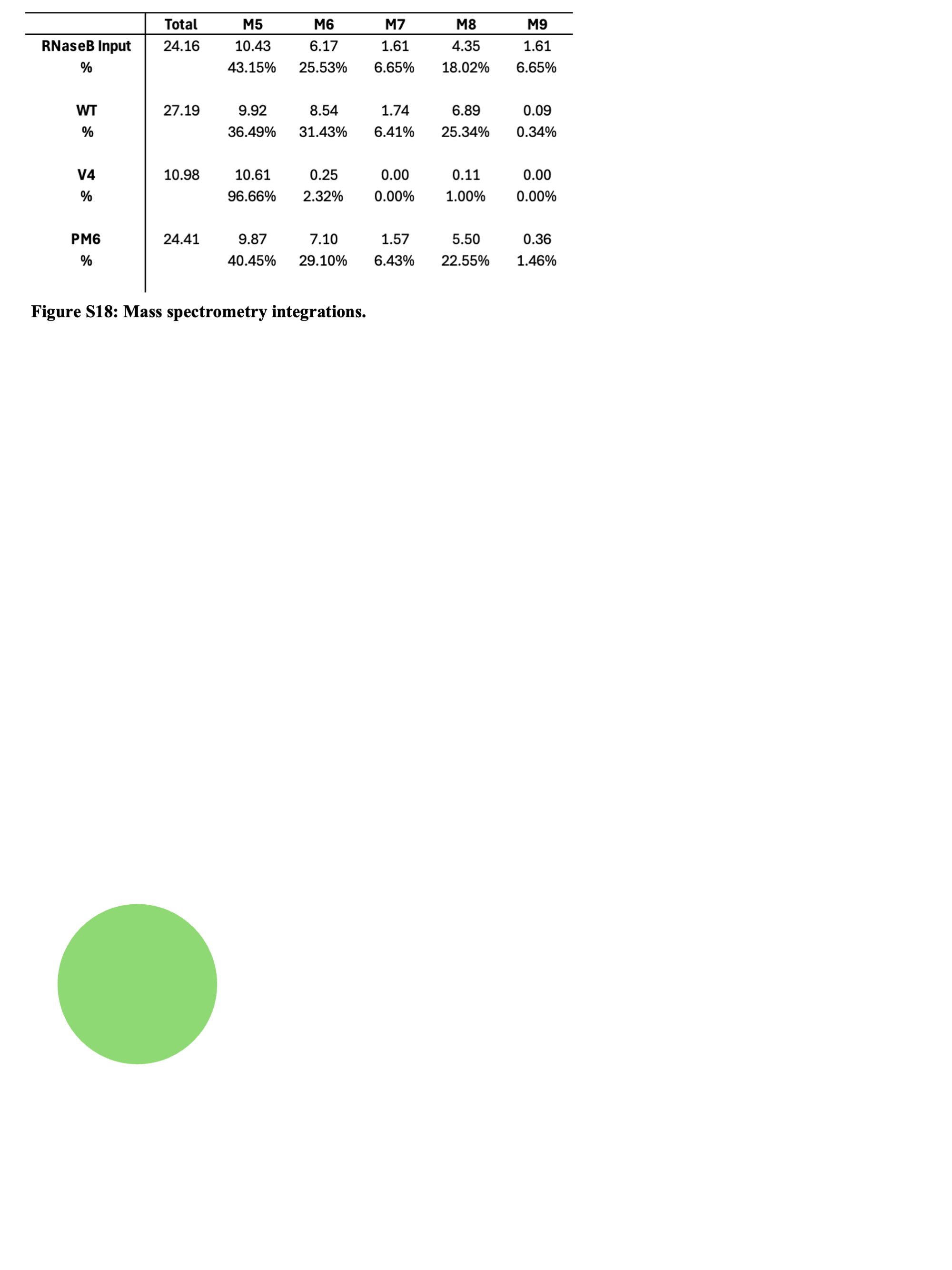

**Supplemental Table 4. Mass spectrometry integrations from RNase B pulldowns.**
